# Bayesian neural networks enable inference of complex phylodynamic processes

**DOI:** 10.64898/2026.02.03.703458

**Authors:** Gabriele Marino, Ugnė Stolz, Cecilia Valenzuela Agüí, Tanja Stadler, Daniele Silvestro

## Abstract

Phylogenetic branching patterns carry essential information about population and diversification dynamic processes, including speciation, extinction, and epidemiological transmission. Phylodynamic models offer a rigorous mathematical framework for quantifying these dynamics from phylogenetic trees. Extensions of these models enable the incorporation of external covariates as predictors of phylodynamic parameters, for instance allowing us to link traits or environmental variables with changes in speciation, extinction, or transmission rates. However, the dependencies between predictors and phylodynamic parameters are typically restricted to linear, additive effects and may thus fail to capture complex, potentially non-linear relationships underlying evolutionary dynamics. To address this limitation, we propose a new framework, BELLA, which leverages unsupervised Bayesian neural networks (BNNs) to flexibly model functional relationships between key phylodynamic parameters and a broad set of predictors, including categorical traits, quantitative variables, and time series data. Based on these covariates, the BNN weights are estimated through Markov chain Monte Carlo and can be inferred jointly with the phylogenetic tree topology and branching times, obtained directly from sequence alignment data. Using extensive simulations, we demonstrate that this approach accurately recovers predictor-parameter relationships, mitigates overfitting, and remains robust across both macroevolutionary and epidemiological contexts. By incorporating tools from explainable artificial intelligence, we further show that our framework reliably identifies the most influential predictors and yields interpretable descriptions of their impact on phylodynamic rates. Finally, we apply our method to two empirical analyses: linking SARS-CoV-2 migration dynamics with travel data during its early spread in Europe, and inferring trait and time-dependent speciation and extinction rates in the Cenozoic diversification of platyrrhines. Our unsupervised BNN framework substantially expands the capabilities of phylodynamic inference providing a powerful and flexible approach to model complex macroevolutionary and epidemiological processes.

**Significance statement:** Understanding how species diversify or pathogens spread requires linking phylogenetic trees to biological traits and environmental characteristics. We present BELLA, an unsupervised Bayesian neural network that learns complex, nonlinear links between predictors—such as traits or environmental variables—and key rates such as speciation, extinction, transmission, and migration directly from sequence data—which in turn inform on the phylogenetic relationships. BELLA does not require any training data, enabling inferences that were previously not tractable. Across simulated epidemics and macroevolutionary histories, BELLA is more accurate than state-of-the-art linear models relating predictors and rates, while remaining interpretable through explainable artificial intelligence. Empirical applications in epidemiology reveal nonlinear effects of travel among countries on the early spread of SARS-CoV-2 in Europe, while a macroevolutionary application shows selective extinction events in the diversifcation history of New World monkeys. The BELLA framework expands the reach of phylodynamics in biology and public health.

## Introduction

Time-calibrated phylogenetic trees are central to understanding the dynamic processes of evolution and populations on macroevolutionary and epidemiological time scales. Such trees are typically inferred from alignments of DNA sequences and morphological traits, with temporal calibration achieved through molecular clock models informed by node-age constraints or time-stamped data, for example fossils of extinct taxa or viral sequences collected through time [1]. The distribution of branching times and lineage durations in these trees provides key information about the underlying population dynamics, which are commonly modeled using birth–death processes [2, 3, 4]. Decades of work in this area have produced a rich class of probabilistic frameworks—collectively known as phylodynamic models—that describe how lineages diversify, spread, and change over time within a phylogenetic context [5, 6, 7].

Phylodynamic modeling has been extensively applied across both epidemiology and macroevolution research. In epidemiology, phylodynamic models are used to infer key parameters that characterize the spatial and temporal development of pathogen outbreaks, such as transmission, migration, recovery, and sampling rates, or the basic reproduction number, *R*_0_, which quantifies the average number of secondary infections caused by an infected individual in a fully susceptible population. These parameters have become instrumental in anticipating the spread of pathogens including SARS-CoV-2, Ebola, Zika, and HIV [8, 9, 10, 11, 12] with application in public health policies [13]. In macroevolution, phylodynamic approaches are one of the pillars for our understanding of how speciation and extinction processes shaped the evolution of biodiversity [14, 6, 15]. Applications of these models have for instance revealed an inverse latitudinal gradient in marine fish speciation rates [16] and consistent diversification dynamics driving the rise and fall of multiple plant and animal clades [17].

Methodological advances in phylodynamics led to models and algorithms that allow us to infer rate variation through time [18], for instance modeling changes of a pathogen effective reproduction number [19] or shifts in speciation and extinction rates [20, 21]. Other models allow for the estimation of rate variations among branches of a phylogenetic tree [22, 23].

Building upon these methods, several models have been developed to explicitly test hypotheses about the impact of biological and environmental factors on population dynamics. Diversity-dependent models, for example, allow researchers to test whether species richness is constrained by ecological limits imposed by finite resources or competition [24, 25]. Trait-dependent diversification models allow us to measure the effect of phenotypic traits on speciation and extinction rates, helping to identify evolutionary key-innovations and providing a mechanistic link between organismal traits and lineage dynamics [26, 27, 28]. Similarly, in epidemiology, multi-type birth–death models enable the estimation of distinct parameters for different subpopulations within a system and quantify the effect of migration among geographic areas on the epidemiological dynamics [29, 30, 31]. Finally, several models have been developed to modulate rate variation as a (generally linear or exponential) function of continuous variables. This has been used in macroevolution to test the impact of environmental variables, for instance global climate change, on species diversification [32, 33]. In epidemiology, generalized linear models (GLMs) have been employed to model the effect of covariates such as host mobility or population density on migration rates between subpopulations [34, 35, 36].

Although these models have greatly improved our ability to quantify population dynamics, existing approaches still face several important limitations. First, they generally rely on overly simple functions— such as piecewise-constant, linear, or exponential forms—to describe rate variation, choices driven more by mathematical convenience than by biological plausibility. Second, most frameworks are either exploratory tools aimed at detecting patterns of rate change, or they are tailored to a single type of predictor, such as an environmental time series or a discrete or continuous trait. Third, while generalized linear models (GLMs) can incorporate multiple predictors, their fixed functional structure restricts them from capturing complex, nonlinear relationships or interactions among variables.

To address these issues, we introduce a novel Bayesian neural network–based framework, extending recent advances in the application of such frameworks to evolutionary studies of fossil data [37]. Neural networks, a class of graphical models, have gained increasing attention due to their capacity as universal approximators, capable of modeling arbitrarily complex functions when provided with sufficient parameters [38]. Typically, they are employed in supervised learning, where mappings between inputs and outputs are inferred from training data. However, in phylodynamic contexts, generating comprehensive training datasets is impractical due to the vast diversity of possible population dynamic scenarios and to limitations in our ability to simulate realistic synthetic datasets [39]. Instead, we propose an unsupervised Bayesian neural network framework for phylodynamic analysis that allows key phylodynamic quantities—such as speciation, extinction, and migration rates, and effective reproduction numbers—to vary both over time and across lineages in response to temporal trends and diverse predictor variables, including traits and environmental time series. Within this framework, phylodynamic parameters are estimated as functions of the predictors, where these functions are defined by neural network weights that are sampled using Markov chain Monte Carlo. This approach eliminates the need for external training datasets and avoids any *a priori* specification of how predictors relate to rate variation.

We demonstrate our framework through extensive simulations spanning a wide range of population dynamic scenarios, from simple constant-rate models to complex nonlinear patterns of rate variation through time and across lineages. We also adapt tools from explainable artificial intelligence (AI) to identify which predictors meaningfully influence epidemiological or macroevolutionary dynamics, providing both accurate inference and interpretability. Finally, we apply our approach to two empirical datasets: estimating migration rates underlying the early spread of COVID-19 in Europe using viral sequence alignments and mobility data, and inferring speciation and extinction dynamics across living and extinct platyrrhines, the New World monkeys.

## Results

### An unsupervised Bayesian neural network model for phylodynamics

We implemented an unsupervised Bayesian neural network to model the variation of phylodynamic parameters as a function of one or multiple predictors, which we termed *Bayesian Evolutionary Layered Learning Architectures* (BELLA, Fig. 1). The BELLA model operates within the phylogenetic inference framework BEAST 2 [40] that can jointly infer tree topology, branch lengths, the parameters of evolutionary substitution models, as well as phylodynamic parameters directly from alignments of molecular and morphological data. The model applies similarly to epidemiology, where the focal phylodynamic parameters may include transmission, recovery, and migration rates [41], and to macroevolution, where speciation, extinction, and fossilization rates describe species diversification and sampling within a fossilized birth–death (FBD) process [42]. The predictors used as input to the neural network include time itself or other time series—such as those describing environmental variation—as well as lineage-specific features, for example phenotypic traits in macroevolutionary processes or population mobility data in epidemiological contexts.

**Figure 1:**
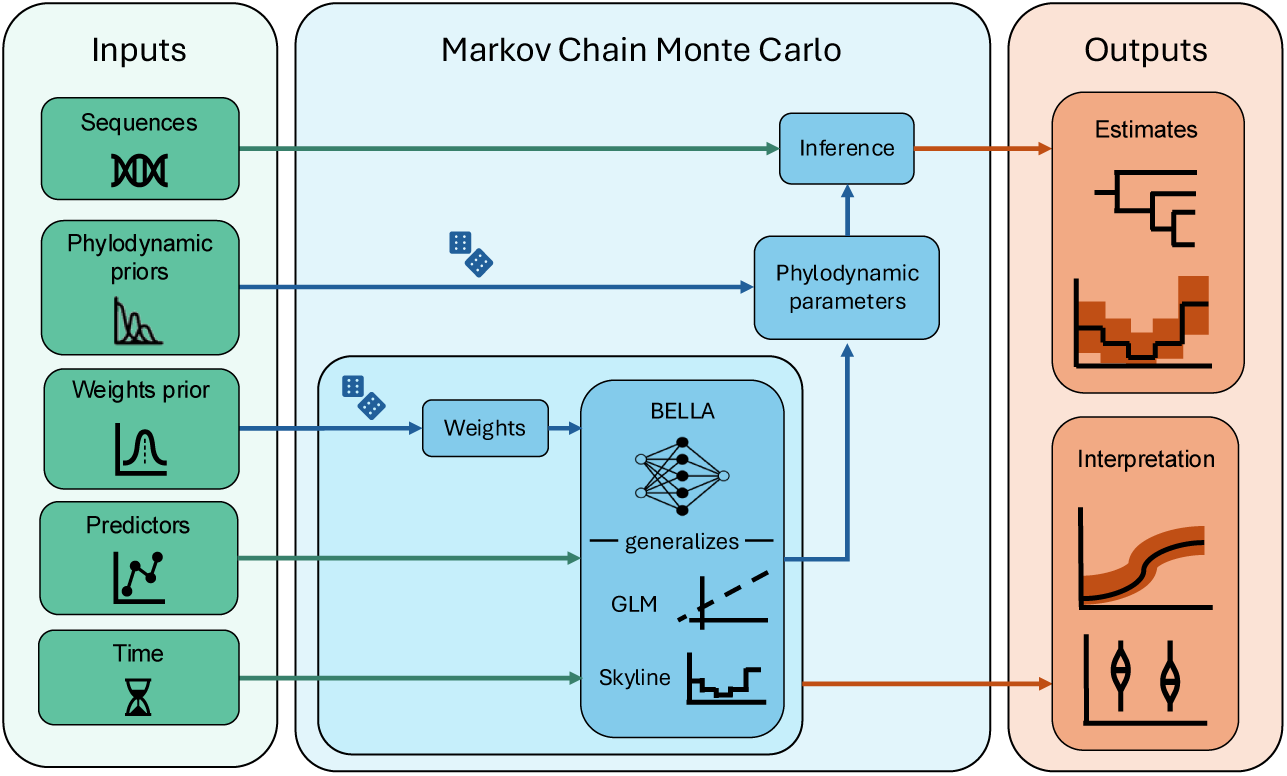
Schematic representation of the proposed *Bayesian evolutionary layered learning architectures* (BELLA) phylodynamic framework. The input data include time-stamped sequencing data (or a fixed, time-calibrated phylogenetic tree) along with predictor variables, such as traits, environmental covariates, or time series. In addition, prior information can be provided for phylodynamic parameters not determined by predictor variables, as well as for the neural network weights. The BELLA model flexibly maps predictors to key phylodynamic parameters—such as speciation, extinction, transmission, or migration rates—allowing for complex, nonlinear relationships that generalize classical generalized linear models (GLMs) and skyline-type approaches. All parameters, including neural network weights, are estimated jointly within a Markov chain Monte Carlo framework. The resulting posterior distributions enable both inference of population dynamics and interpretation of predictor effects using explainable artificial intelligence tools.

In the implemented framework, we employ a phylodynamic birth–death model [43] to describe the branching of the tree, in which the phylodynamic parameters are the birth, death, migration, and sampling rates, determined by predictors through time and across lineages. While our framework is described here in the context of the birth–death model, the same concepts can also be applied to other phylodynamic approaches, such as the coalescent [44].

As in standard phylodynamic analyses, the BELLA model relies on unsupervised Bayesian inference to sample the parameters from their posterior probability distribution using a Markov chain Monte Carlo (MCMC) algorithm. Yet, in contrast to most implementations, the BELLA model does not sample the parameters of the phylodynamic process directly; instead, it samples the weights of the neural network, from which such phylodynamic parameters are obtained. Owing to its representational flexibility, the neural network serves as a proxy for the potentially complex mapping between input predictors and phylodynamic parameters, enabling the model to capture nonlinear, non-monotonic responses and interactions among predictors.

### Performance of the BELLA model

We assessed the performance of our model through analyses of a diverse set of simulated datasets, and showed that the BELLA model reliably achieved accurate results in both epidemiological and macroevolutionary contexts. We tested different architectures of fully connected neural networks with two hidden layers and different numbers of nodes (see *Methods* for more details), including a small network (3 and 2 nodes, respectively), an intermediate one (16 and 8 nodes), and a larger one (32 and 16 nodes). We hereafter report results from the intermediate network, which on average performed best across the datasets considered, and provide all results in the *Supplementary Information*.

The simulation scenarios included a varied suite of predictor–parameter relationships, encompassing constant, linear, and nonlinear dependencies applied to the reproduction number and migration rates (epidemiology; Fig. S1a–d), as well as to speciation and extinction rates (macroevolution; Fig. S1e–h). We evaluated the relative performance of the BELLA models by analyzing all simulated datasets under state-of-the-art alternatives, including “predictor-agnostic” models, in which rate variation is modeled using parameters that vary independently across time or lineages (for instance, piecewise-constant rates through time) [18, 41], and generalized linear models (GLMs) that modulate rate variation through linear correlations with the predictors [34, 35, 36].

We first assessed the performance of the BELLA model in scenarios where the simulated predictor-parameter relationship was constant (Fig. S1a,e). Although time and a random time series were included as predictors, neither introduced bias into the inference, and the model successfully recovered the constant rates with high accuracy and precision (Figs. S2a, S3a). The mean absolute errors (MAE) and the average widths of the 95% credible intervals were comparable to or smaller than those obtained from GLMs for both reproduction numbers and speciation and extinction rates (Tabs. S1, S3). Coverage exceeded 0.95, indicating that the true parameter values were consistently captured within the posterior distributions (Tab. S2). By contrast, the predictor-agnostic model – which estimated independent phylodynamic parameters across the ten time bins – generally showed higher error and lower coverage, with particularly poor estimates in early time bins (Figs. S2a, S3a). This pattern likely reflects overfitting due to the lack of regularization or model selection procedures.

Simulation scenarios with linear relationships between input predictors and phylodynamic rates demonstrated that the BELLA model can accurately capture these linear trends without showing signs of overfitting (Figs. S1b,f). Overall, BELLA performed comparably to—or in some cases slightly better than—the GLM in terms of MAE, coverage and average width of the 95% credible intervals (Figs. S2b, S3b; Tabs. S1–S3). For example, in a macroevolutionary scenario where speciation rates increased linearly and extinction rates declined over time (Fig. S1b), BELLA achieved an MAE of 0.012 and a coverage of 0.97. In contrast, the GLM exhibited a higher MAE (0.020) and slightly lower coverage (0.91), although its 95% credible intervals were narrower on average (0.12) than those of BELLA (0.16).

We also evaluated BELLA under a macroevolutionary scenario where both speciation (*λ*) and extinction (*µ*) rates varied through time and across lineages as a function of a binary trait evolving along the tree (Fig. S1h). In this setting with interacting predictors, BELLA achieved the lowest MAE (0.013), outperforming the GLM (0.021), and provided better coverage (0.98 versus the GLM’s 0.91), although with slightly wider average 95% CI width (0.24 compared to GLM’s 0.16).

Finally, we assessed the performance of BELLA in scenarios with nonlinear predictor-parameter relationships. These included epidemiological simulations with non-monotonic reproduction numbers over time (Fig. S1c) and logistic patterns of migration rates (Fig. S1d), as well as macroevolutionary simulations featuring declining speciation rates and spiking extinction rates (Fig. S1g). In these complex settings, BELLA successfully captured the non-linear rate patterns (Figs. 2, S2c, S3c), achieving higher accuracy than both predictor-agnostic models and GLMs, as reflected by lower MAE (Table S1). For example, migration rates estimated with BELLA (MAE = 0.0036) were 1.9 times more accurate than those from the GLM (MAE = 0.0070) and 16.1 times more accurate than those from the predictor-agnostic model (MAE = 0.058). Similarly, the accuracy of estimated speciation and extinction rates improved 4- to 5-fold under BELLA compared with the alternative approaches. Coverage remained high across the epidemiological simulations (0.96–0.98), representing a slight improvement over predictor-agnostic models (0.90–0.96) and a substantial improvement over GLMs, whose coverage dropped to 0.61–0.75 due to their limited flexibility in capturing rate variation (Tab. S2). In the macroevolutionary simulations, BELLA’s coverage decreased to 0.73 for extinction rates, reflecting a slight underestimation of the single-time-bin peak (Fig. S3c). Finally, the average width of the 95% credible intervals was again comparable between BELLA and the GLM model, and substantially smaller than that obtained using the predictor-agnostic model (Tab. S3).

**Figure 2:**
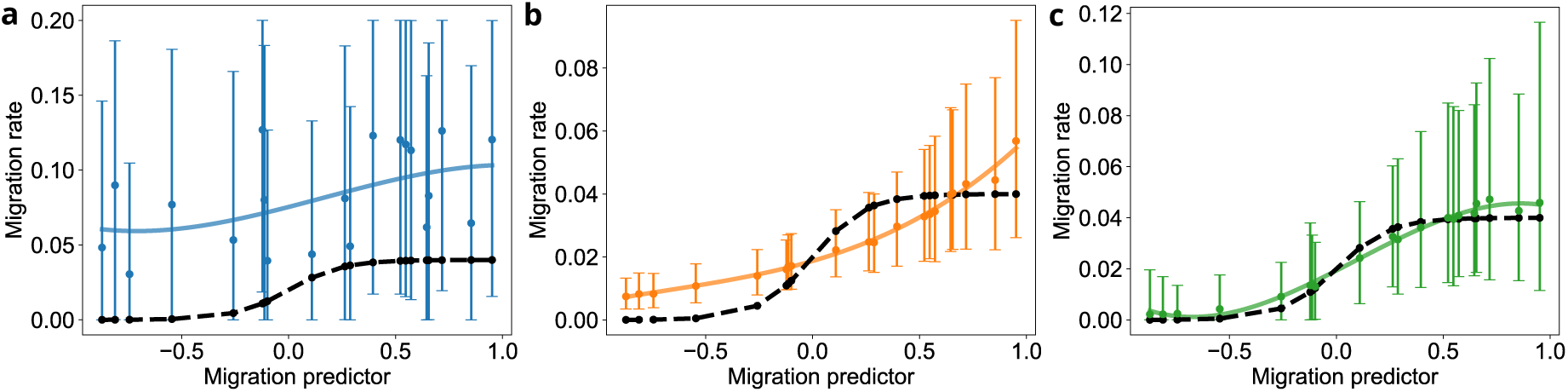
Results for a simulation scenario involving the prediction of 20 migration rates among all pairs of 5 populations (Epi-4). Migration rates were modeled based on a logistic function of an input predictor. Each panel shows the median estimates and 95% credible intervals averaged across 100 simulated phylogenies for: (a) a predictor-agnostic model (blue), (b) a GLM model (orange), and (c) the BELLA model (green). Solid lines indicate a 1-D smoothing spline fit, dashed lines show the true migration rates.

### Interpretation of the BELLA model using explainable artificial intelligence

To interpret the influence of predictors on population dynamics, we implemented algorithms adapted from explainable artificial intelligence (xAI). Specifically, we visualized the inferred effects of predictors on rate variation using partial dependence plots (PDPs), which extract the marginal effect of an individual predictor while averaging over all others. We also computed the Shapley values, to quantify the average contribution of each predictor to the model output, thus allowing us to rank the predictor importance. Together, these complementary metrics quantify both the direction and magnitude of each predictor’s impact on the inferred phylodynamic parameters.

We evaluated our interpretation framework on a macroevolutionary scenario in which speciation (*λ*) and extinction (*µ*) rates varied both over time and among species, depending on a binary trait evolving across the phylogeny (Fig. S1h). The analysis also included a random time series and an additional binary trait, neither of which influenced speciation or extinction. Partial dependence plots (PDPs) successfully captured the effects of the relevant predictors (time and trait 1) by illustrating their marginal impact on BELLA’s output, while indicating negligible effects for the irrelevant predictors (random time series and trait 2) (Fig. 3a, b). PDPs with respect to time revealed that the model correctly associated time with decreasing speciation and increasing extinction rates, whereas the random time series showed minimal variation (Fig. 3a). Similarly, the first binary trait was correctly linked to shifts in rates, while the second trait had no discernible effect (Fig. 3b). These results demonstrate that BELLA effectively captured both the temporal trend and the influence of the relevant trait, while remaining largely insensitive to irrelevant predictors. This conclusion is further supported by Shapley analysis (Fig. 3c), which identified time and the relevant trait as the most influential features, with the other predictors contributing minimally. Finally, the median predicted speciation rates across all species classes (Fig. 3d) confirm that BELLA accurately recovered the distinct phylodynamics incorporated in the simulation.

**Figure 3:**
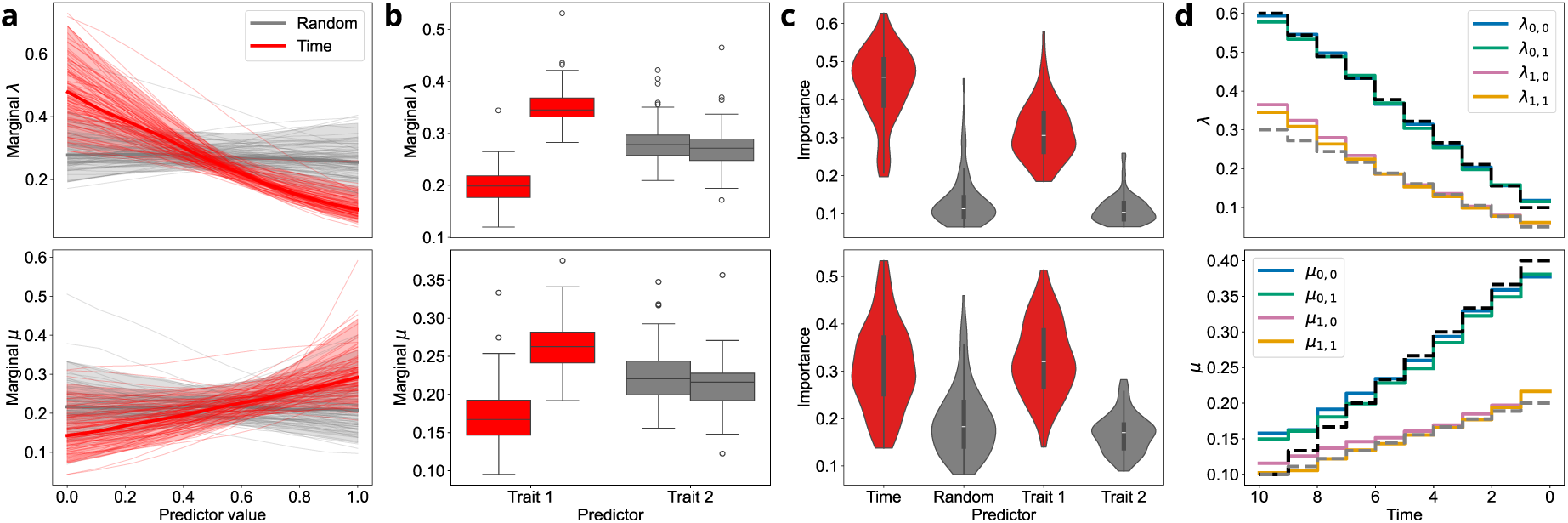
Results for a simulation scenario featuring time-varying speciation (*λ*) and extinction (*µ*) rates across four groups of species as a function of a binary trait evolving across the tree (FBD-4). The analysis includes four predictors: time (relevant, shown in red), a random continuous predictor (irrelevant, gray), and two binary traits: trait 1 (relevant) and trait 2 (irrelevant). For 100 simulated phylogenies, the panels show: (a) the distributions of partial dependence plots (PDPs) for the two continuous predictors (time and random); (b) the distributions of PDPs for the two binary traits, with left and right boxes corresponding to trait values 0 and 1, respectively; (c) the distributions of Shapley feature importances; the (d) median predicted rates through time. In the panels, the top row corresponds to the speciation rate (*λ*) and the bottom row to the extinction rate (*µ*).

### Results on empirical data

#### Speciation and extinction dynamics in New World monkeys

We applied our BELLA framework to explore macroevolutionary dynamics in the New World monkeys, to estimate their speciation and extinction rates from a set of 100 phylogenetic trees of 87 extant and 34 extinct species from a previous Bayesian analysis [45]. We used time (binned into geological stages) and body mass (discretized into four categories) as predictors of rate variation (Fig. 4).

**Figure 4:**
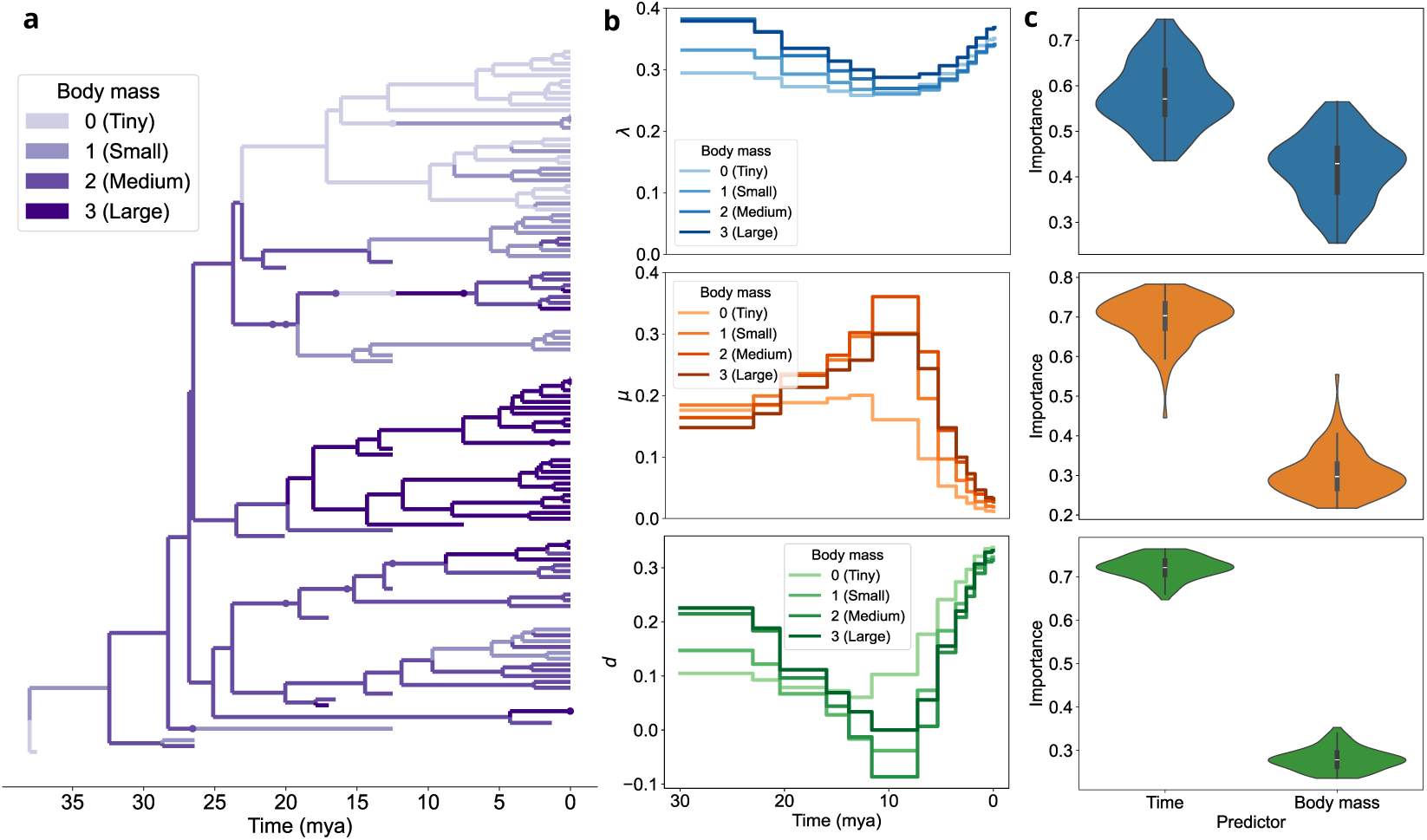
Diversification analysis of New World monkeys (Platyrrhines) using the BELLA framework. The analysis estimates time-varying speciation rates (*λ*, top row), extinction rates (*µ*, middle row), and net diversification rates (*d* = *λ* − *µ*, bottom row) as functions of time and a discretized body-mass trait evolving along the phylogeny. (a) The evolution of the discretized body-mass trait across the maximum clade credibility tree. (b) The median posterior estimates across time and body-mass categories for each macroevolutionary rate (full distributions are presented in Fig. S4). (c) Shapley feature importance values for each predictor and macroevolutionary rate.

Overall, the model captures a mixed influence of time and body mass on the evolutionary dynamics. The speciation rate (*λ*) remains relatively stable across both predictors, but shows a ca. 20–30% drop during the early Miocene (between 23 and 10 Ma) across lineages with body-mass above 500 g (categories 1, 2 and 3). In contrast, speciation rates in the smallest taxa (body-mass category 0) are only marginally affected by this event, exhibiting a decrease of approximately 11%. Speciation rates increase starting from the late Miocene and through the Recent, converging to similar values across all body-mass categories.

Extinction rates (*µ*) exhibit a stronger variation and interaction between time and body mass. Larger-bodied species (body-mass categories 1, 2 and 3) experienced a 1.6–2.2 fold spike in extinction around 10 Ma whereas smaller-sized species were inferred to be largely unaffected by this event. Extinction rates rapidly decreased across all body-mass categories starting in the late Miocene, a trend that continued through the Recent.

The diversification rates (*d* = *λ* − *µ*) reflect these trends and interaction between time and body mass. Larger-bodied species underwent an event of negative diversification rate that peaked around 10 Ma, while smaller-bodied species maintained a relatively stable rate. After this event, all body-mass groups exhibit an upward trend in diversification, leading to positive rates and implying a recent accumulation of diversity in the clade.

The Shapley-based analysis of feature importance revealed similar contributions of time and body mass to the speciation rate. In contrast, for the extinction and diversification rates, time received substantially greater importance, reflecting the dominance of temporally structured macroevolutionary dynamics—such as clade-wide extinction and diversification pulses—over trait-driven effects. As a consequence, body mass was assigned lower overall importance for extinction and diversification, although its influence is still evident in how it separates the spiking extinction and diversification rates of larger-bodied groups while leaving those of the smallest species relatively stable.

#### Phylogeographic reconstruction of early SARS-CoV-2 spread in Europe

In all previously presented applications of the BELLA model, the phylogenetic tree topology was fixed. However, our approach can be seamlessly incorporated into a full phylogenetic inference pipeline in which the tree is jointly estimated from multiple sequence alignment data. To illustrate this in a real-world setting, we applied BELLA to estimate SARS-CoV-2 migration rates across countries during the early spread of SARS-CoV-2 in China and Europe, using 123 viral genomes sampled between the onset of the outbreak and 8 March 2020.

We modeled migration rates as a function of the number of flights between countries, normalized by the population size of the source country, following previous analyses [36], and compared the BELLA results with those obtained using a GLM. The phylogenetic trees inferred by the two approaches were generally consistent (normalized Robinson–Foulds distance = 0.64 [46]; Fig. S8) with both trees sharing the same overall structure with only minor topological differences. The results also showed agreement in the reconstructed geographic spread of the outbreak, with both analyses identifying Italy as the most probable location of the most recent common ancestor of the European clade (Fig. 5a; Fig. S9). We examined how BELLA and the GLM link migration rates to the normalized number of flights used as predictor through partial dependence plots (PDPs, Fig. 5c). BELLA inferred a less steep dependency, that flattens slightly at low and high predictor values.

**Figure 5:**
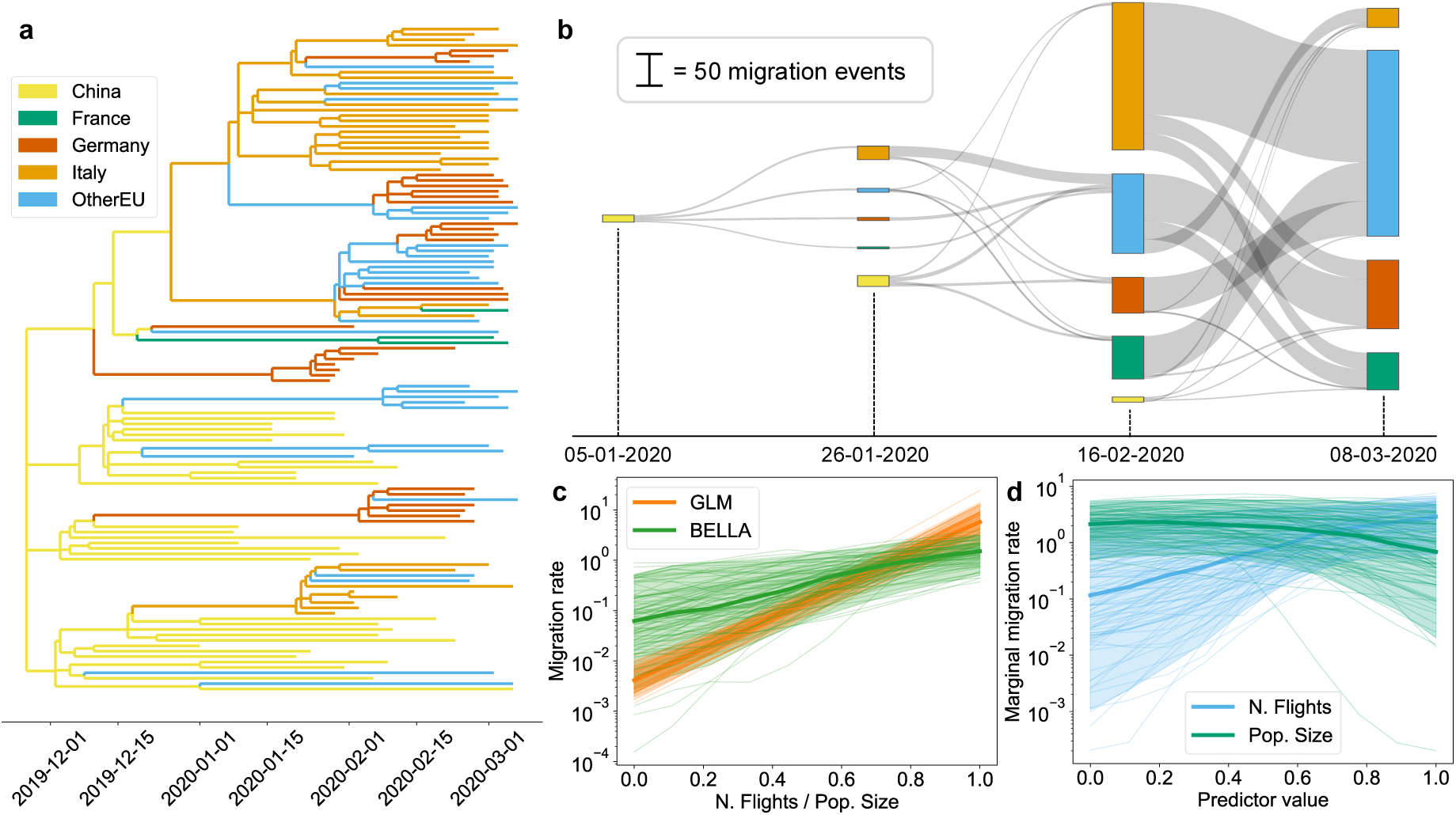
Application of the BELLA model to estimate migration rates across China and European countries at the onset of the COVID-19 pandemic. (a) The inferred transmission maximum clade credibility tree. (b) Median estimated migration fluxes between countries (full distributions are presented in Figs. S12–S14) (c) Partial dependence plots (PDPs) for BELLA and a generalized linear model (GLM) comparing the effect of the normalized number of flights on predicted marginal migration rates. (d) PDPs illustrating the disentangled effects of the number of flights and the source country population size on predicted marginal migration rates.

To further disentangle the contribution of individual predictors, we ran an additional analysis using flight counts and population size as separate predictors in BELLA. The resulting PDPs (Fig. 5d) indicate that population size has little effect except at very large values, where a slight inverse effect emerges. In contrast, flight volume shows a nonlinear response, with a threshold-like pattern: a steep increase at low values that gradually tapers off at higher values.

Finally, we applied a phylogenetic particle filtering algorithm [47] to sample trajectories from the posterior distribution of the population history given the genetic data. This approach allowed us to estimate migration fluxes between countries over the study period. The estimated fluxes were consistent between BELLA and the GLM, with Italy inferred as the principal exporter of viral lineages and Germany as the main importer during this phase of the epidemic (Fig. 5b; Fig. S10, Figs. S12–S14). Similarly, both models yielded consistent estimates of the distribution of introduction dates from China to Europe, with the majority of introductions occurring around mid-January 2020 (Fig. S11).

## Discussion

### Modeling population dynamics with unsupervised Bayesian neural networks

The increasing amount of sequencing data enables phylodynamic inference to be a cornerstone for the quantification of population and diversification dynamic parameters. In macroevolutionary studies, these parameters are typically expressed as speciation and extinction rates [14, 6, 15], whereas in epidemiological settings they describe quantities such as migration and transmission rates and effective reproduction numbers [8, 9, 10, 11, 12].

Although evolutionary dynamics are likely influenced by multiple factors, most existing models are limited to the effects of single (or few) predictors, such as time-varying rates [20, 48, 32], trait-dependent diversification [26], or the combination of the two [41]. Predictor values are typically binned into a small number of discrete classes; for example, time may determine birth and death rates in a piecewise-constant fashion, independently across time bins. In such models, increasing the number of classes provides additional flexibility but simultaneously inflates the number of parameters and their associated uncertainty, ultimately leading to uninformative parameter estimates. Some models allow for multiple predictors or maintain a continuous state space [28, 49, 34, 35, 36]. However, they typically rely on simplifying assumptions, such as constraining rate variation to be monotonic and additive, and often neglect interactions between covariates.

The Bayesian evolutionary layered learning architecture (BELLA) framework relaxes many of these assumptions through an unsupervised neural network. The approach allows us to jointly infer the effects of multiple predictors in modeling rates dynamics through time and across lineages. Our simulations demonstrate that BELLA performs robustly across a broad spectrum of epidemiological and macroevolutionary scenarios, including those featuring nonmonotonic effects and interacting covariates. Our approach reliably recovered a wide range of predictor–parameter relationships with high accuracy and precision, while providing well-calibrated uncertainty estimates, consistently matching or outperforming competing phylodynamic methods, including predictor-agnostic inference and generalized linear models. Performance gains were particularly pronounced in scenarios with nonlinear predictor effects and in those involving interacting predictors. Importantly, even in scenarios with simple constant or linear relationships, BELLA matched or exceeded the performance of the existing GLM framework in both accuracy and precision.

BELLA integrates two complementary sources of information: hypothesis-driven, biologically-motivated predictors—such as traits that may influence speciation or travel patterns that inform migration—and forms of rate variation that make no assumptions about underlying mechanisms, including generic temporal changes. This combination is critical; indeed, many birth–death models adopt a constant-rate process as the null expectation, an assumption that is known to bias inference and generate spurious signals of trait or environmental dependence [26, 50, 28, 51, 52]. Although BELLA’s complex parameterization structure makes traditional hypothesis-testing tools (e.g., Bayes factors for individual predictors) impractical, we demonstrate that methods from explainable AI can be effectively used to highlight which predictors exert the strongest influence on inferred phylodynamics. This aligns with recent studies showing that Shapley-based approaches can recover drivers of survival during mass extinction events [53] and identify both trait-dependent diversification and human-induced extinctions in proboscideans [37]. While such techniques cannot formally reject predictors, they do provide reliable estimates of their relative contributions. In our simulations—including those with interacting traits—BELLA consistently recovered accurate measures of predictor influence. However, as with any high-dimensional regression framework, strong correlations among predictors or time series can limit interpretability. Careful selection of ecologically, biologically, or environmentally relevant variables therefore remains essential when designing BELLA analyses [52].

Recent work has shown that neural networks can recover speciation, extinction, transmission, and recovery rates from fossil records or phylogenies, but these studies typically operate within a supervised learning paradigm [54, 55, 56, 57, 58, 59]. In those approaches, researchers produce large collections of simulated datasets generated under known macroevolutionary or epidemiological processes, extract features from those simulations, and then train neural networks to map those features back to the underlying rate parameters. Once trained, the models are applied to real data to estimate rate dynamics. While this strategy has proven effective for relatively simple scenarios—such as models with limited trait or time dependence—it becomes increasingly impractical when moving toward more intricate settings like those addressed by BELLA, where rates may vary in complex, high-dimensional ways. Constructing a sufficiently broad training library to cover such possibilities would require an enormous and often infeasible simulation effort. In contrast, our framework avoids the entire supervised-learning pipeline. Rather than relying on pre-labeled simulated data, BELLA learns directly from the empirical dataset by evaluating its likelihood under a birth–death process. This unsupervised approach makes it possible to infer how predictors shape diversification or transmission dynamics without specifying a priori which kinds of rate variation are allowed or how predictors might interact.

Neural networks offer considerable freedom in how predictor variables can influence diversification or transmission rates, but that flexibility comes with the risk of introducing far more parameters than the data can actually inform. In standard frequentist applications—especially in supervised learning where networks are trained on large labeled datasets—this excess capacity must be tightly controlled through explicit regularization strategies such as dropout, weight penalties, or early stopping based on validation performance [60]. Without such safeguards, many of the network’s weights simply absorb noise rather than signal. Bayesian neural networks, however, naturally guard against this problem because the prior distributions imposed on the weights act as intrinsic regularizers [61, 37]. This property is key to the performance of our approach, which accurately recovered also constant-rate processes.

A related consideration concerns the size of the neural network itself. In supervised settings, it is common to use large architectures—sometimes extremely wide or deep—because abundant simulated or empirical training data can support such complexity, and the goal is to maximize predictive power.

In our framework, the network plays a different role: it serves as a flexible but generic mapping from predictors to rate parameters, not as a prediction engine trained on labeled examples. Because of this, we employed comparatively compact architectures. Although we found that smaller architectures performed better in constant scenarios and larger ones in nonlinear settings or when many predictors were used, these differences were not decisive. Overall, BELLA delivered consistent inference quality, indicating low sensitivity to the specifics of the architecture. This stability is important, as it shows that BELLA does not require extensive or scenario-specific hyperparameter tuning to produce reliable results, enhancing its practicality and general applicability for phylodynamic analyses.

A further advantage of adopting a Bayesian neural network is that uncertainty is naturally quantified. Because the weights are treated as random variables and jointly explored through their posterior distribution, the framework yields full posterior distributions for population dynamic rates. Importantly, although the BELLA model contains many parameters, all of them are informed by the entire dataset. For example, increasing the number of time bins when inferring time-dependent rates does not introduce additional independent parameters, and uncertainty does not become extreme in periods with sparse data. In contrast, piecewise-constant models, which treat rates in each time bin as independent parameters, can exhibit large uncertainty in data-sparse regions such as early time bins. In this respect, BELLA is particularly advantageous in time bins with little or no data, as information can be shared across the full dataset through common network weights.

### Diversification history of the New World Monkeys

BELLA analyses place the origin of Platyrrhines around 40 Ma in a small-bodied ancestor weighing less than 500 g (category 0, *Tiny*, in our discretized body-mass framework). This estimate is consistent with interpretations of the earliest fossil evidence for the clade [62]. Subsequently, the clade diversified into multiple larger-bodied lineages, some of which exceeded 3000 g (category 3, *Large*) by the early Miocene (ca. 20 Ma; Fig. 4a). This reconstruction corroborates previous inferences based on continuous models of trait evolution [45].

The evolution of platyrrhines is characterized by substantial variation in speciation and extinction rates, which our BELLA model attributes to heterogeneities through time and among body-mass categories. Temporal effects were strongest for extinction, with rates peaking in the late Miocene (11–7 Ma), consistent with previous findings [45]. This extinction peak is likely linked to the combined effects of global cooling following the Middle Miocene Climatic Optimum and the aridification of southern South America associated with Andean uplift and rain-shadow effects [63]. Larger taxa exhibited overall higher speciation and extinction rates, meaning higher turnover, a pattern widely observed across fossil mammals [64]. Partial dependence plots further revealed that large-bodied lineages were disproportionately affected by the late Miocene extinction event. Together, these results point to a non-additive interaction between temporal and trait-dependent effects on extinction, a pattern that would be difficult to detect using alternative phylodynamic models.

The sustained decline in extinction rates, together with a moderate increase in speciation rates, leads to strongly positive net diversification toward the present, in agreement with previous findings based on simpler time-dependent phylodynamic models applied to extant-species phylogenies [65]. Speciation and extinction rates through time are also remarkably consistent with paleodiversity estimates inferred from an independently compiled dataset of fossil occurrences and derived using different methodologies and assumptions [66]. Notably, that analysis identified a rapid decline in predicted species richness during the late Miocene, followed by a rebound—dynamics that our results here attribute to a simultaneous increase in speciation and decrease in extinction.

While the credible intervals around the marginal rates through time remain wide (Fig. S5), they are substantially narrower than those obtained in a previous FBD analysis based on the same phylogenetic trees, which assumed a piecewise-constant model with independent rate parameters across time bins regularized by a Brownian process prior [45]. In particular, the upper bounds of the rates (here approximately 0.8 for speciation and 0.6 for extinction events per lineage per million years) are roughly tenfold lower than the highest values estimated previously. This highlights BELLA’s ability to yield more tightly constrained uncertainty intervals than predictor-agnostic piecewise-constant models, a pattern also observed in simulation analyses (Figs. S2, S3; Tab. S3).

### Early European phylogeography of SARS-CoV-2

Following the emergence of SARS-CoV-2 in China at the end of 2019, Europe experienced large-scale COVID-19 outbreaks in early 2020. Here we investigate early global spread through phylodynamic analyses. Building on prior studies that incorporate international air-travel data as a predictor of migration rates, we employ a flexible unsupervised Bayesian neural network to describe the relationship between travel volumes and migration, instead of the GLM framework used previously [36]. While we did not perform a formal model comparison between BELLA and the GLM, the two approaches produced similar results in terms of phylogeographic reconstruction, and both converged to nearly identical phylogenetic likelihoods (Fig. S7).

Because the GLM is limited in capturing interactions between predictors, careful feature engineering– such as normalizing flight volumes by the source-country population–is important [36]. BELLA’s flexibility, instead, allows it to account for interactions between predictors, and when repeating the analysis with the two predictors disentangled, we converged to very similar results, suggesting that the key patterns of migration inferred from the data are robust to different model specifications, and that BELLA can accommodate predictor interactions without extensive manual preprocessing.

Based on the BELLA output, our analysis employs a phylogenetic particle filtering algorithm to sample epidemic trajectories from the posterior distribution of the population history [47, 48], thereby extending earlier work [67]. The inferred transmission history indicates that initial introductions into Europe occurred around mid-January 2020, earlier than suggested by confirmed case reports S11. This timing aligns both with previous phylodynamic reconstructions [36] and with retrospective serological evidence suggesting that SARS-CoV-2 may have been circulating in parts of Europe as early as December 2019 [68, 69]. Differences arise, however, when examining the inferred migration fluxes between countries (Fig. S14). The GLM tends to produce distributions that are more constrained and peaked, giving a narrower representation of plausible migration fluxes. BELLA, in contrast, produces wider, flatter distributions, providing a more nuanced view of migration patterns and capturing the uncertainty typical of early-stage phylodynamic epidemiological reconstructions.

### Conclusions

Inferring population and diversification dynamic processes from biological data is essential for understanding the forces shaping biodiversity and the spread of pathogens across space and time. This task is inherently challenging because available data are incomplete: only a small fraction of ancient taxa leave a fossil record, many extant species remain unsampled, and in epidemic settings only a subset of infected individuals is typically sequenced. Phylodynamic methods provide a principled framework for reconstructing past dynamics from such incomplete data. However, as models become more complex, parameter uncertainty can increase to the point that inferences lose interpretability, and mechanistic links between key processes—such as speciation, extinction, and migration—and environmental or biotic drivers remain difficult to quantify. Our BELLA approach enables accurate, data-driven reconstruction of complex phylodynamic patterns while controlling overfitting and retaining interpretability through explainable AI techniques.

We envision this framework enabling robust phylodynamic inference across a wide range of application domains, including macroevolution and epidemiology, as demonstrated here, as well as areas such as single-cell biology and linguistics where phylodynamic tools are increasingly applied. Applications naturally include phylogeography, which in our formulation falls within the broader phylodynamic framework. More generally, this approach opens the door to incorporating mechanistic relationships between polulation dynamic parameters and diverse predictors, ranging from life-history traits to environmental variables. Together, these developments provide a foundation for more general and mechanistically grounded phylodynamic inference, advancing our understanding of the processes that shape population dynamics and evolution.

## Methods

### Modeling phylodynamic rates with a neural network

Phylodynamic inference seeks to quantify the population dynamic and sampling processes that give rise to an observed phylogenetic tree *T* ; we refer to the model of this process as the *phylodynamic model*. In macroevolutionary applications, the relevant parameters typically include speciation, extinction, and fossilization rates, whereas in epidemiological settings they may include transmission, recovery, and sampling rates. A widely used model for population dynamics is the birth–death model, parameterized by a birth rate *λ*, a death rate *µ*, and a sampling rate *ψ* [29]. More elaborate birth–death models may additionally incorporate migration processes and type-dependent rates [41]. While this work focuses on phylodynamic inference under the birth–death framework, the concepts developed here are also applicable to the coalescent framework [44].

The posterior distribution over the phylodynamic parameters is given by

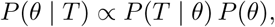

where *θ* = (*λ, µ, …*) denotes the set of parameters of the birth–death model. The phylodynamic like-lihood *P* (*T* | *θ*) specifies the probability of observing the tree topology and branch lengths under the assumed birth–death process. Phylodynamic inference therefore aims to identify the parameter values that most plausibly generated a given phylogenetic tree, while naturally accounting for uncertainty through the posterior distribution. In practice, phylogenetic trees are typically not observed directly but are themselves inferred from empirical data, such as aligned molecular sequences or morphological characters. Consequently, the phylodynamic process is commonly used as a tree prior within a joint Bayesian framework in which the phylogeny is estimated simultaneously with the phylodynamic parameters.

Most phylodynamic models directly estimate the parameters of a birth–death process, such as speciation and extinction rates in macroevolution, or migration rates and reproduction numbers in epidemiology. Alternative approaches instead model these rates as functions of external predictors. In generalized linear model (GLM)–based formulations, phylodynamic parameters are given by a (tipically logarithmic) link function applied to a linear combination of predictors. The inferred parameters are the intercept and regression coefficients of this relationship [34, 35, 36]. Here, we model variation in phylodynamic rates as a function of predictors using a feedforward neural network.

A feedforward neural network (FFNN) is a flexible function approximator capable of capturing complex, nonlinear relationships between inputs and outputs [38]. An FFNN consists of an input layer, one or more hidden layers, and an output layer. Each layer applies a linear transformation followed by a nonlinear activation function. Formally, for layer *l* with input vector **x**^(*l*−1)^, weight matrix **W**^(*l*)^, and bias vector **b**^(*l*)^, the output is given by

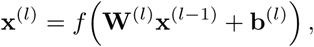

where *f* (·) denotes a nonlinear activation function, such as the rectified linear unit (ReLU) or the sigmoid function. By stacking multiple layers, an FFNN can approximate arbitrary continuous functions, enabling it to learn complex mappings from predictor variables to phylodynamic rates.

In standard applications, FFNNs are used in supervised learning settings, where the network is trained to map inputs to observed target values by minimizing a loss function, such as mean squared error for regression or cross-entropy for classification. In this context, the network weights and biases are treated as parameters that are optimized during training. In contrast, BELLA adopts an unsupervised use of FFNNs to integrate external, non-phylogenetic data into phylodynamic analyses. The FFNN maps these input predictors to a subset of the phylodynamic parameters, denoted *θ*_FFNN_, while priors are placed on the network weights **W** and biases **b**. The remaining parameters, *θ*_base_, are not predicted by the FFNN and are instead sampled directly from their standard priors. This separation allows the model to exploit external covariates for parameters that can be informed by additional data, while retaining classical Bayesian sampling for the remaining parameters.

To ensure that the FFNN produces parameter values within biologically or mathematically meaningful ranges, the choice of output activation functions can be tailored to the specific parameter being modeled. For instance, parameters that are inherently bounded between zero and one, such as probabilities, can be mapped through a sigmoid or logistic transformation. More generally, sigmoidal-like functions with user-specified upper and lower bounds allow parameters to be constrained to finite intervals, while exponential or softplus activations can enforce strict positivity. By aligning the output activation function with the natural support of each parameter, the model avoids generating implausible values and improves the efficiency of posterior exploration.

As in standard phylodynamic analyses, Markov chain Monte Carlo (MCMC) [70] is used to jointly sample from the posterior

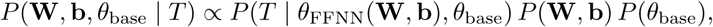

with operators acting on both the network parameters and the standard prior-sampled parameters. At each MCMC iteration, new values for **W** and **b** are proposed and used by BELLA to map the input predictors to the phylodynamic rates *θ*_FFNN_, which are then combined with *θ*_base_ to evaluate the phylodynamic likelihood. When phylodynamic rates vary across lineages, the predictor values associated with individual branches are typically unknown. For instance, the geographic location or trait state of an ancestral branch is rarely observed. In such cases, the FFNN maps predictor values to lineage-specific rates for each possible ancestral state, and the likelihood is computed by integrating over all these possibilities [41]. This procedure naturally propagates uncertainty about ancestral predictor values through the neural network and into the posterior distribution, providing a fully Bayesian integration of external data into phylodynamic inference.

### Evaluation metrics

To compare the performance of the different inference approaches (predictor-agnostic, GLM, and BELLA), we computed the posterior median of each target parameter from every MCMC run, along with the corresponding 95% credible intervals (CIs). Model performance within each scenario was evaluated based on three complementary metrics: the mean absolute error (MAE) between the posterior median estimates and the true parameter values across the 100 simulated trees, the average width of the 95% CIs, and the empirical coverage of these intervals.

Formally, let ***θ****_p_* = (*θ_p_*_1_ *, …, θ_pK_*) denote the vector of true values for parameter *p*, where *K* is the number of elements in the vector (e.g., time bins for piecewise-constant time-varying parameters or population pairs for migration rates), and let 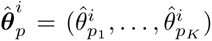 be the corresponding posterior median estimates obtained from tree *i*, with *i* = 1*, …, N* and *N* = 100 simulated trees per scenario. The mean absolute error (MAE) for each parameter and each tree was computed as

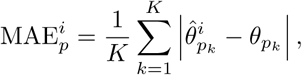

and the overall MAE for a given parameter in a given scenario was then obtained by averaging across all trees:

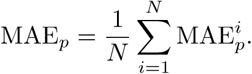

The width of the 95% credible interval (CIW) for element *k* of parameter *p* in tree *i* was defined as

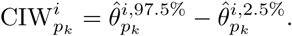

The average CIW for parameter *p* in a given scenario was then obtained by first averaging over all elements of the parameter vector for each tree and then across all trees:

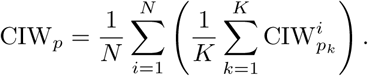

Coverage was defined as the proportion of true parameter values contained within their respective 95% credible intervals. For each element *k* of parameter *p* in tree *i*, let

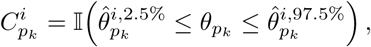

where I(·) denotes the indicator function. The overall coverage for parameter *p* was then obtained by averaging over all elements and all trees:

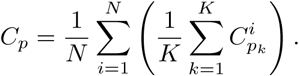

The mean absolute error quantifies overall estimation accuracy, the average width of the 95% credible interval reflects the precision of posterior uncertainty, and the coverage measures the calibration of the credible intervals. Together, these three criteria provide a comprehensive assessment of both point estimates and uncertainty quantification across models and scenarios.

To assess the computational efficiency and sampling performance of each inference framework, we computed the mean effective sample size per hour. For each model and scenario, we first estimated the effective sample size (ESS) [71] of all inference targets obtained from the posterior samples. The ESS quantifies the number of effectively independent draws from the posterior, accounting for autocorrelation in the MCMC chain. The mean ESS across all parameters was then divided by the total wall-clock time (in hours) required for the corresponding inference run. This procedure was repeated across all replicate phylogenetic trees, yielding the mean ESS/hour averaged over all replicates for each scenario and model combination, allowing for direct comparison of computation efficiency between models.

### Interpretability analysis

To better understand and interpret the behavior of FFNNs, which are often regarded as “black-box” models due to their complex and non-linear structure, we employed two explainable AI (xAI) techniques [72]: partial dependence plots (PDPs) [73] and Shapley values [74]. These methods allow us to quantify the influence of individual predictors on model outputs and to visualize their effects across the predictor space. In particular, we used them to assess the robustness of BELLA to irrelevant input predictors, as well as to gain general insights into how the models leverage relevant features in the inference task.

Partial dependence plots (PDPs) illustrate the average effect of a single predictor *x_S_* (or a subset of predictors) on the model’s output, marginalizing over the distribution of all other features *x_C_*. Formally, for a prediction function *f* (**x**), the partial dependence of the feature subset *S* is defined as

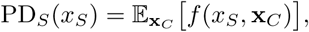

where *C* denotes the complement of *S*, and the expectation is taken with respect to the marginal distribution of the remaining features, *p*(**x***_C_*).

PDPs are particularly useful because they provide an interpretable, model-agnostic visualization of how an input feature influences the prediction on average, even in the presence of complex nonlinear interactions captured by models such as FFNNs. If the predictor is relevant, the PDP typically exhibits a structured relationship (e.g., monotonic trend, threshold effect, or nonlinear curve). Conversely, if the predictor is irrelevant, the PDP should remain approximately flat, indicating that varying this feature does not systematically alter the model output. This makes PDPs well-suited for diagnosing spurious dependencies and detecting overfitting when irrelevant predictors are included in the input space. However, PDPs also have important limitations, as they implicitly assume independence between features when marginalizing, which may lead to misleading plots if predictors are correlated.

Shapley addresses these issues by offering a game-theoretic framework for feature attribution. For a model *f* and input instance **x**, the Shapley value for feature *i* is:

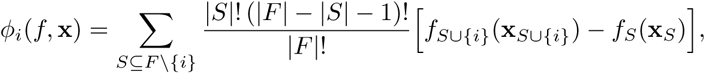

where *F* is the full feature set and *S* a subset excluding *i*, and the term *f_S_*(**x***_S_*) is defined through a conditional expectation:

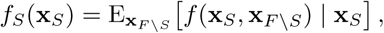

that is, the model output averaged over the distribution of the missing features given the known ones.

Shapley values provide local, prediction-level explanations, but in practice, feature importances are often summarized by averaging the absolute Shapley values across all instances in the input dataset:

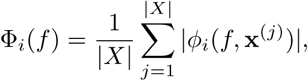

where |*X*| is the number of samples. This provides a global quantification of each feature’s contribution, complementing the local and marginal insights offered by PDPs, and enables a more comprehensive assessment of how predictors influence the network outputs across different conditions.

In our study, each FFNN has multiple configurations due to the weights sampled during the MCMC process. To compute PDPs and Shapley values, we first subsampled 100 MCMC samples per run, yielding 100 distinct FFNN models per tree. For each input predictor, we computed the PDP values for each model, and then aggregated them into a single PDP per tree by taking their median. Similarly, Shapley values were computed for each of the 100 subsampled FFNN models, then aggregated by computing the median feature importance per tree, and finally marginalized so that the feature contributions per tree sum to 1.

### Simulation scenarios

We benchmarked our approach on eight simulation scenarios, grouped into two classes of multi-type birth–death models [29]: epidemiological processes (with removal after sampling) and fossilized birth– death (FBD) processes [42] (with ancestor sampling and complete present-day sampling) (Fig. S1).

All epidemiological scenarios share the same become-uninfectious rate *δ* = *µ* + *ψ* = 0.07, the same sampling probability upon removal 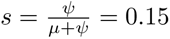, and the same simulation time *T* = 250. Scenarios Epi-1, Epi-2, and Epi-3 involve a single homogeneous population, with the inference task being to estimate a time-varying reproduction number, *R_t_* = *λ/*(*µ* + *ψ*), assumed piecewise constant, using time as the predictor. *R_t_* is constant in scenario Epi-1, linearly decreasing in scenario Epi-2, and non-monotonic in scenario Epi-3. Scenario Epi-4 is characterized by five populations, with constant reproduction numbers for each of them (respectively, 0.8, 1.0, 1.2, 1.4 and 1.6), and the task is to infer the 20 pairwise migration rates using a continuous covariate as migration predictor. To generate these rates, the migration predictors were first sampled from a uniform distribution u(−1, 1) and then mapped into target migration rates using a logistic function.

All FBD scenarios assume the same constant sampling rate through time *ψ* = 0.2 and simulation time *T* = 35, and the inference task is to estimate piecewise-constant speciation and extinction rates across 10 time bins. Predictors are given by time and, for scenario FBD-4, by two additional binary traits. Scenario FBD-1 assumes constant speciation and extinction rates; scenario FBD-2 features a linearly decreasing speciation rate and a linearly increasing extinction rate; scenario FBD-3 models nonlinear decreases in the speciation rate, while the extinction rate exhibits a sharp upward spike in the second half of the simulation. Scenario FBD-4 involves four different groups of species, distinguished by the combination of values of two binary traits. However, the second of these traits does not affect population dynamics, meaning that the four populations can be further grouped into two pairs that share identical speciation and extinction rates. For each pair, the speciation rate decreases linearly while the extinction rate increases linearly.

To assess the robustness of our approach to misspecifications in the predictor set—particularly with respect to the inclusion of irrelevant predictors—we incorporated an additional randomly generated predictor into all simulation studies. This predictor was constructed randomly and independently of the inference target, and resampled for each tree. In scenario Epi-4, it consisted of a uniformly distributed random value in [1, −1] for each of the 20 population pairs, whereas for all other scenarios, it was defined as a random walk with standard normal steps across the time bins.

For each scenario, we simulated 100 phylogenetic trees with 200 to 500 tips. While BELLA can be fully integrated within the tree inference environment in BEAST, we here fixed the trees to reduce the substantially higher computational cost associated with joint tree inference. An example of BELLA combined with tree inference is presented in one of our empirical analyses on the early spread of SARS-CoV-2 in Europe (see below).

We performed MCMC inference on each tree using the BDMM-Prime package [41] implemented in BEAST 2 [40] coupled with different inference approaches: (i) a predictor-agnostic model that estimates each parameter independently (e.g., inferring the value of a piecewise-constant rate for each time bin without shared structure), (ii) a GLM implementation [75] that estimates an intercept and growth factors defining an exponential mapping between predictors and phylodynamic parameters, and (iii) three BELLA architectures. The BELLA networks had hidden layer sizes of (3, 2), (16, 8), and (32, 16) neurons, with ReLU activations in the hidden layers and a logistic activation in the output layer. For analyses requiring multiple parameters (e.g., speciation and extinction rates in FBD scenarios), a separate network was used for each target parameter, with the output layer consisting of a single neuron. The input layer size varied according to the number of predictors in each experiment, and all predictors were normalized to the range [0, 1] prior to inference.

For the predictor-agnostic model and GLMs, we employed uniform priors for the phylodynamic parameters: U(1, 5) for reproduction numbers, U(0, 0.05) for migration rates, and U(0, 2) for speciation and extinction rates. We used standard normal priors N (0, 1) for the weights of the neural networks in BELLA, and for the intercept and growth factors in the GLMs. For BELLA, we also constrained the output phylodynamic parameters through a logistic output activation to match the bounds of the corresponding uniform priors.

All MCMC runs employed chains of 10,000,000 steps. For all analyses, the first 10% of samples were discarded as burn-in, and convergence was assessed by verifying that the effective sample size for each parameter exceeded 200.

### Diversification analysis of New World monkeys

We analyzed a sample of 100 dated phylogenetic trees encompassing 121 species of the Platyrrhine clade, of which 87 are extant and 34 are extinct. These trees represent a posterior sample from a previously published analysis of 55 genes for extant species, combined with multiple topological constraints for extinct taxa based on taxonomic assessments [45]. In our analysis, we investigated how speciation and extinction rates vary as functions of time and body mass, both of which have been hypothesized to capture heterogeneity in the diversification dynamics of the clade [76, 45]. Time was incorporated using geological stages as discrete bins, with the Late Pleistocene and Holocene merged into a single bin (0.129–0 Ma) to avoid intervals of disproportionately short duration. The model also included sampling rates through time, which were assumed to follow a piecewise-constant process with independent rates for each geological epoch (Eocene, Miocene, and Pliocene); the Pleistocene and Holocene were similarly merged given the short duration of the latter.

Because the BDMM-Prime package [41] does not currently support continuous traits, we discretized body-mass values into four ordinal categories, resulting in subsets of species of comparable size: category 0 (tiny), up to 500 g (21 taxa); category 1 (small), 500–1000 g (26 taxa); category 2 (medium), 1000–3000 g (34 taxa); and category 3 (large), above 3000 g (32 taxa). Body-mass values were unavailable for eight taxa and were therefore treated as missing data in the analysis. To reflect the ordinal nature of this trait, we specified a transition matrix that allowed direct transitions only between neighboring categories.

After setting the sampling fraction at the present to 0.44 (as in the original study [45]), we ran 10,000,000 million MCMC iterations to sample the parameters of the model. These included the transition rates among adjacent body-mass categories, the sampling rates through time and the weights of the BELLA framework, mapping time and body mass to speciation and extinction rates.

For BELLA, we used a neural network with two hidden layers comprising 16 and 8 neurons, respectively, corresponding to the architecture that performed best on average in our simulation studies. Input predictors were normalized to the interval [0, 1], and a standard normal prior N (0, 1) was assigned to the network weights. We employed ReLU activations in the hidden layers and a logistic activation in the output layer to constrain the predicted rates to the positive range [0, 5]. Priors on the sampling rate through time and on the transition rates between body-mass groups were specified over the same range.

For all analyses, the first 10% of samples were discarded as burn-in, and convergence was assessed by verifying that the effective sample size for each parameter exceeded 200. To aggregate estimates across the 100 phylogenies into a single representative tree, we computed a maximum clade credibility (MCC) tree using the TreeAnnotator software bundled with BEAST 2 [40].

### Phylogeographic analysis of early SARS-CoV-2 spread in Europe

We analyzed the early spread of COVID-19 across Europe using an open access dataset comprising 123 SARS-CoV-2 full viral genomes sampled between the start of the pandemic and 8 March 2020.

The sequences were downloaded from GenBank [77] on 16 January 2026 via LAPIS [78]. We retained sequences with a Nextclade QC overall score below 100 to maximize data availability while excluding those flagged as *bad* [79]. This score accounts for missing data (fewer than 3,000 nucleotides), mixed sites, private mutation clusters, and premature stop codons. Finally, we masked the first 100 and last 200 nucleotides, as well as additional flagged sites, following the Nextstrain ncov pipeline [80]. The dataset included 30 genomes randomly selected from China (the origin of the pandemic), 30 from Germany, and 30 from Italy. For France, three sequences were included: although 20 sequences were available during the study period, 17 belonged to a single hospital cluster. To avoid bias from over-represented transmission chains, only one representative from this cluster was retained. Of the remaining three sequences, two passed quality filters. The remaining European countries were combined into a composite “Other European” deme, from which 30 genomes were randomly selected from all other available European sequences, with country-level sampling probabilities weighted by the number of cases reported by the European Centre for Disease Prevention and Control (ECDC) during the study period.

We used these sequences to jointly infer transmission dynamics and the phylogeny. For the evolution of the sequences along the tree, we assumed an HKY model [81] with empirical base frequencies. Rate heterogeneity among sites was accommodated with four gamma-distributed categories, with the gamma shape parameter assigned an exponential prior (mean 0.5). The transition/transversion ratio (*κ*) was assigned a lognormal prior (mean 1.0, scale 1.25). We further assumed a strict clock with a fixed substitution rate of 8.0 × 10^−4^ substitutions per site per year [67].

For the phylodynamic model, we used the multi-type birth–death model implemented in BDMM-prime [41] combined with BELLA and with a GLM. The model included deme-specific effective reproduction numbers, sampling proportions, and pairwise migration rates [67]. The reproductive number was assumed to remain constant during the analysis period for the European demes, whereas a temporal change was incorporated for China on the date of the Hubei lockdown (23 January 2020). We specified lognormal priors for the origin time of the birth–death process (mean -1, scale 0.2) and for the reproduction numbers (mean 0.8, scale 0.5). Sampling proportions were assigned uniform priors, bounded between the ratio of observed sequences to confirmed cases and one-tenth of that value, reflecting the assumption that the true number of infections ranged from one to ten times the number of confirmed cases.

We modeled migration rates between countries using air travel connectivity as a predictor, considering two parameterizations: one based on flight passenger volumes together with source-country population sizes, and another using passenger volumes normalized by source-country population size. This choice was motivated by repeated evidence that air travel strongly shapes the spatial spread of respiratory viruses, including SARS-CoV-2 [82, 36]. We used flight volume data from the Eurostat transport datasets (avia paexcc [83] and avia paocc [84]) for December 2019 and January–March 2020, together with country population data from the 2020 World Development Indicators [85].

The mapping from input predictors to migration rates was parameterized using a generalized linear model (GLM) and a BELLA model consisting of two hidden layers with 16 and 8 neurons, respectively. We used standard normal priors N (0, 1) for the neural network weights in BELLA and for the growth factors in the GLM. The GLM intercept was assigned a lognormal prior (mean = 0, scale = 1), and BELLA employed a Softplus output activation function. For each analysis, we ran three independent MCMC chains with different random seeds, each consisting of 10,000,000 steps, and verified convergence by confirming that all chains reached the same posterior distributions and that the effective sample size for each parameter exceeded 200. The first 10% of samples were discarded as burn-in. To summarize phylogenetic uncertainty, we computed a maximum clade credibility (MCC) tree using TreeAnnotator from BEAST 2 [40].

To estimate migration fluxes between countries, we sampled multi-type population trajectories using the phylogenetic particle filtering algorithm [47] as implemented in BDMM-Prime [41] from the posterior distribution obtained through the phylodynamic analysis described above. For each sampled tree and set of model parameters, the algorithm simulates an ensemble of possible histories weighted by their agreement with the tree and parameters, from which one history is drawn. Applying this procedure to a subsample of states visited by the MCMC chain yields a set of trajectories representing the posterior distribution of the population history given the genetic data.

### Computational runtime

The increased parameter space associated with neural networks compared to predictor-agnostic or GLM-based formulations make them computationally slower (Tab. S4); overall the runtime difference varied by roughly a factor of 10. As expected, smaller neural networks tend to converge faster in simple settings, whereas larger architectures may be preferable when handling a greater number of predictors. We ran both empirical analyses for 24 hours, by which point all chains had reached a substantial number of effective samples (ESS *>>* 200) for each estimated parameter.

### Implementation and data availability

Our approach is implemented in the BEAST 2 framework [40] as a new package, *Bayesian Evolutionary Layered Learning Architectures* (BELLA), which is publicly available at https://github.com/gabriele-marino/BELLA/. BELLA can be seamlessly combined with a wide range of birth–death population models, sequence evolution models, and the various operators provided in BEAST 2, enabling phylodynamic inference to be performed either jointly with phylogenetic tree estimation—accounting for tree uncertainty—or based on a fixed, known tree.

All code required to reproduce the analyses presented here is available at https://github.com/gabriele-marino/BELLA-companion, including code used to simulate the benchmarking data; the empirical New World monkey phylogenies; the aligned SARS-CoV-2 sequences with GenBank identifiers; the BEAST configuration files used to run inference on both simulated and empirical data; and the post-processing and interpretation pipelines based on partial dependence plots and Shapley feature importance.

## Acknowledgments

The authors thank ETH Zürich for funding. TS and US received funding from the European Research Council (ERC) under the European Union’s Horizon 2020 research and innovation programme grant agreement no. 101001077. BEAST 2 development is partly funded by Swiss Institute of Bioinformatics (SIB). We thank Tim Vaughan for his valuable guidance and support on this work, particularly regarding the use of BDMM-Prime and the BEAST 2 software suite. We also thank Chaoran Chen for his prompt and helpful assistance in collecting publicly available SARS-CoV-2 sequences. Finally, we thank the entire Computational Evolution team at ETH Zürich for continuous discussions and feedback on the project.

## Supplementary Information

### Supplementary Figures

**Figure S1:**
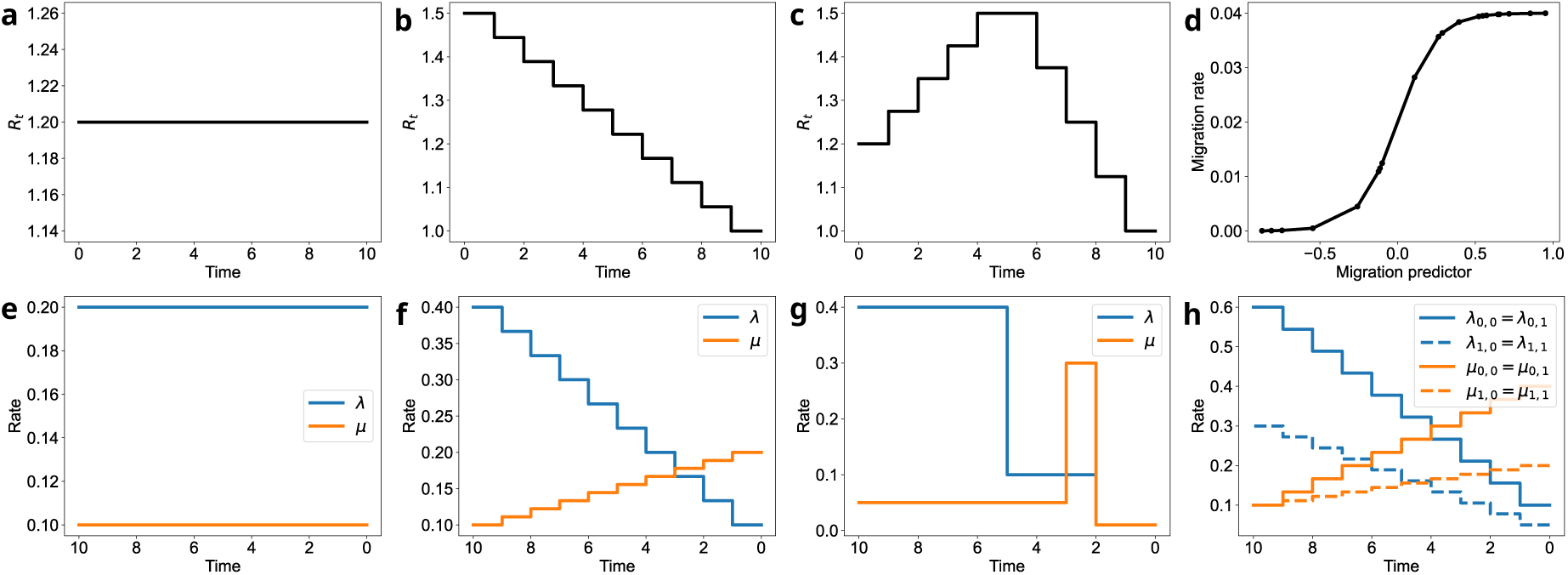
The simulation scenarios considered in this study. Each panel corresponds to a specific scenario and illustrates the relationship between input predictors and the resulting phylodynamic rates. (a) Epi-1: inference of a constant reproduction number using time as a predictor; (b) Epi-2: inference of a linearly decreasing reproduction number using time as a predictor; (c) Epi-3: inference of a non-monotonic reproduction number using time as a predictor; (d) Epi-4: inference of migration rates between five populations predicted by a continuous covariate; (e) FBD-1: inference of constant speciation and extinction rates using time as a predictor; (f) FBD-2: inference of linearly decreasing speciation rate and linearly increasing extinction rate using time as a predictor; (g) FBD-3: inference of a non-linear speciation rate with a spike in the extinction rate using time as a predictor; (h) FBD-4: inference of linearly decreasing speciation rate and linearly increasing extinction rate across four groups of species defined by two binary traits (only one trait affects diversification), using time and trait values as predictors.

**Figure S2:**
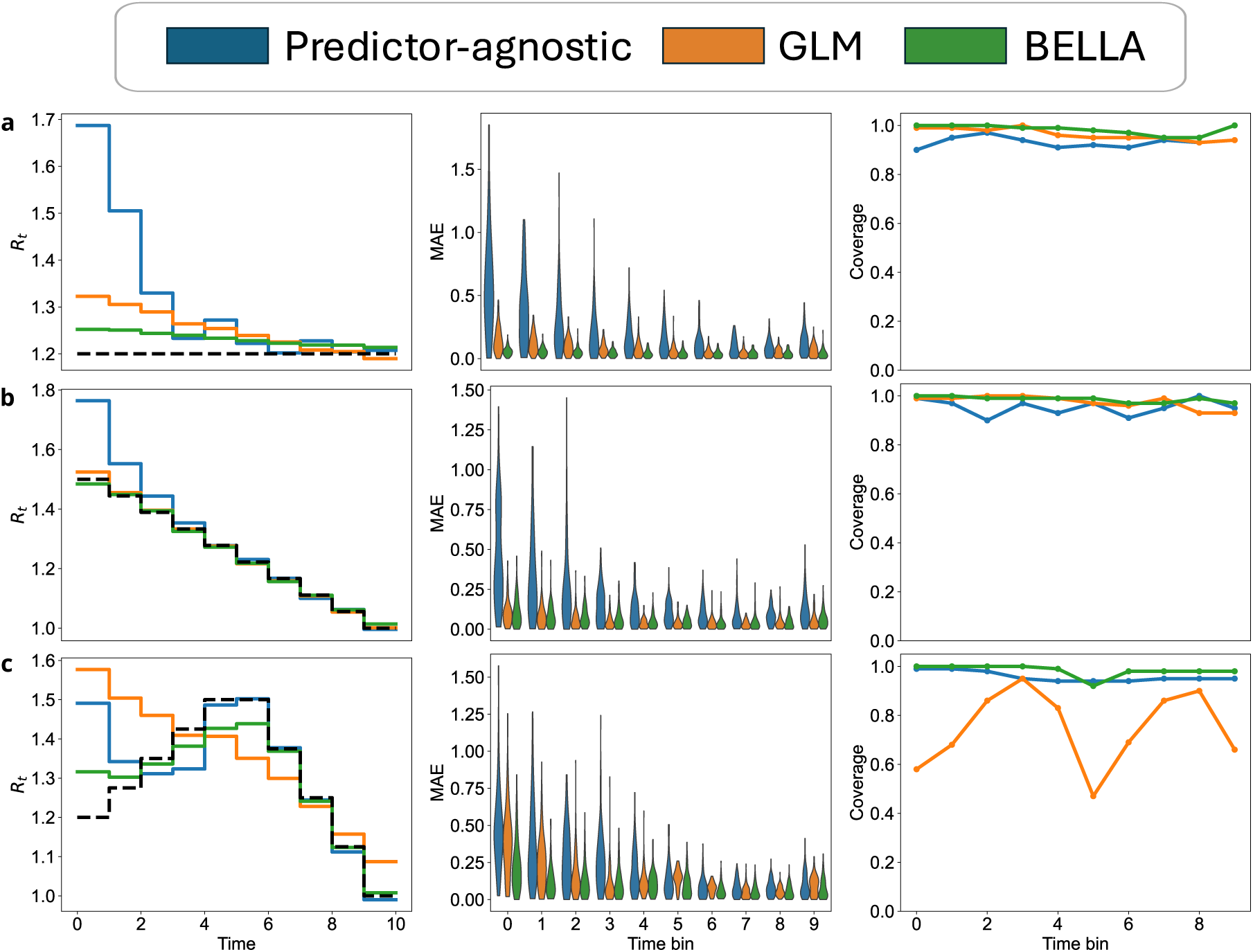
Results for the epidemiological simulation scenarios: (a) Epi-1; (b) Epi-2; (c) Epi-3. For each inference framework— predictor-agnostic (blue), GLM (orange), and BELLA (green)—the panels display, from left to right: the median reproduction number estimates across all replicate trees; the distribution of the mean absolute error (MAE) across trees; and the coverage of the 95% credible intervals.

**Figure S3:**
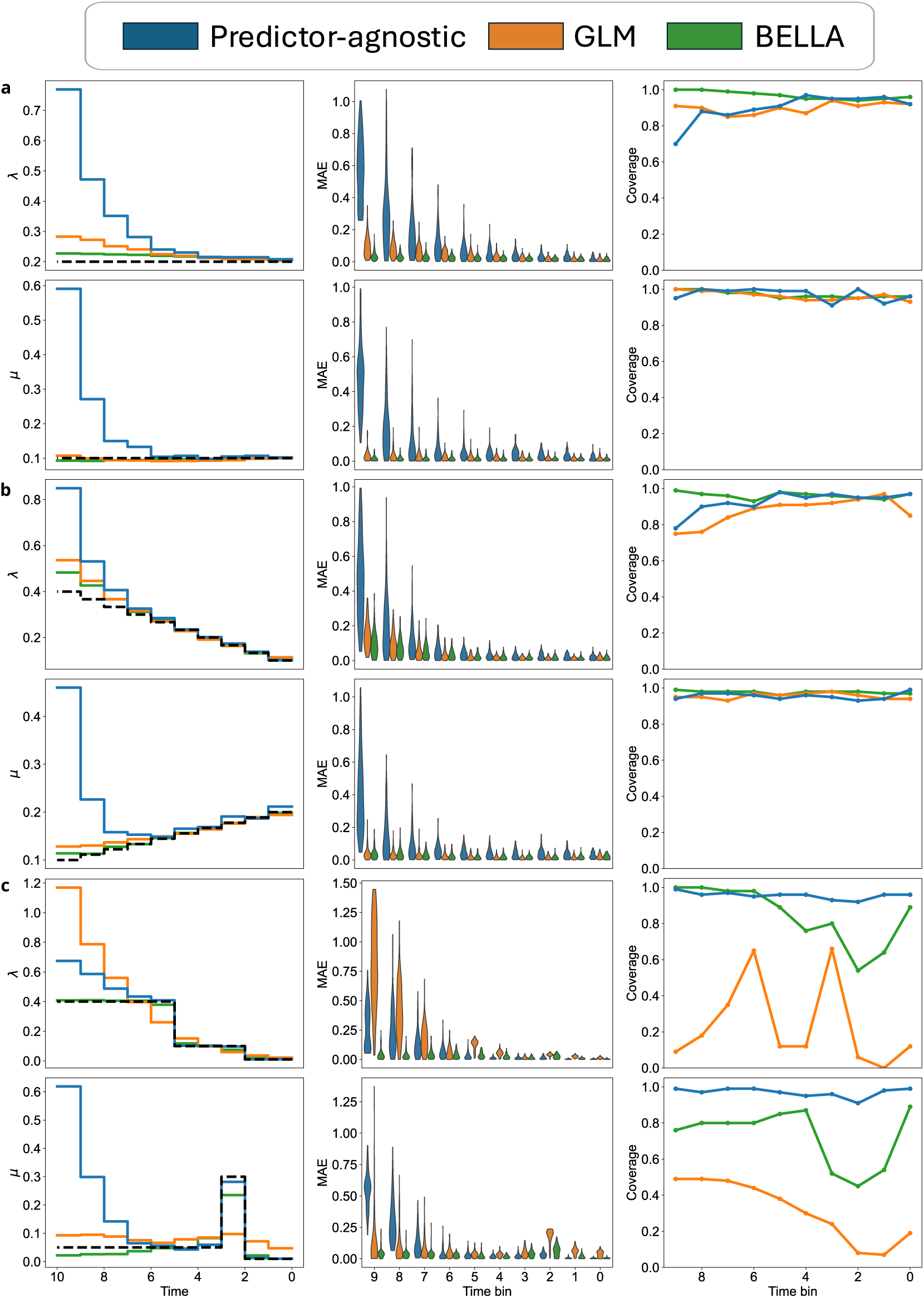
Results for the FBD simulation scenarios: (a) FBD-1; (b) FBD-2; (c) FBD-3. For each inference framework—predictor-agnostic (blue), GLM (orange), and BELLA (green)— and for each inference target (speciation rate *λ*, top row; extinction rate *µ*, bottom row), the panels show, from left to right: the median rate estimates across all replicate trees; the distribution of the mean absolute error (MAE) across trees; and the coverage of the 95% credible intervals.

**Figure S4:**
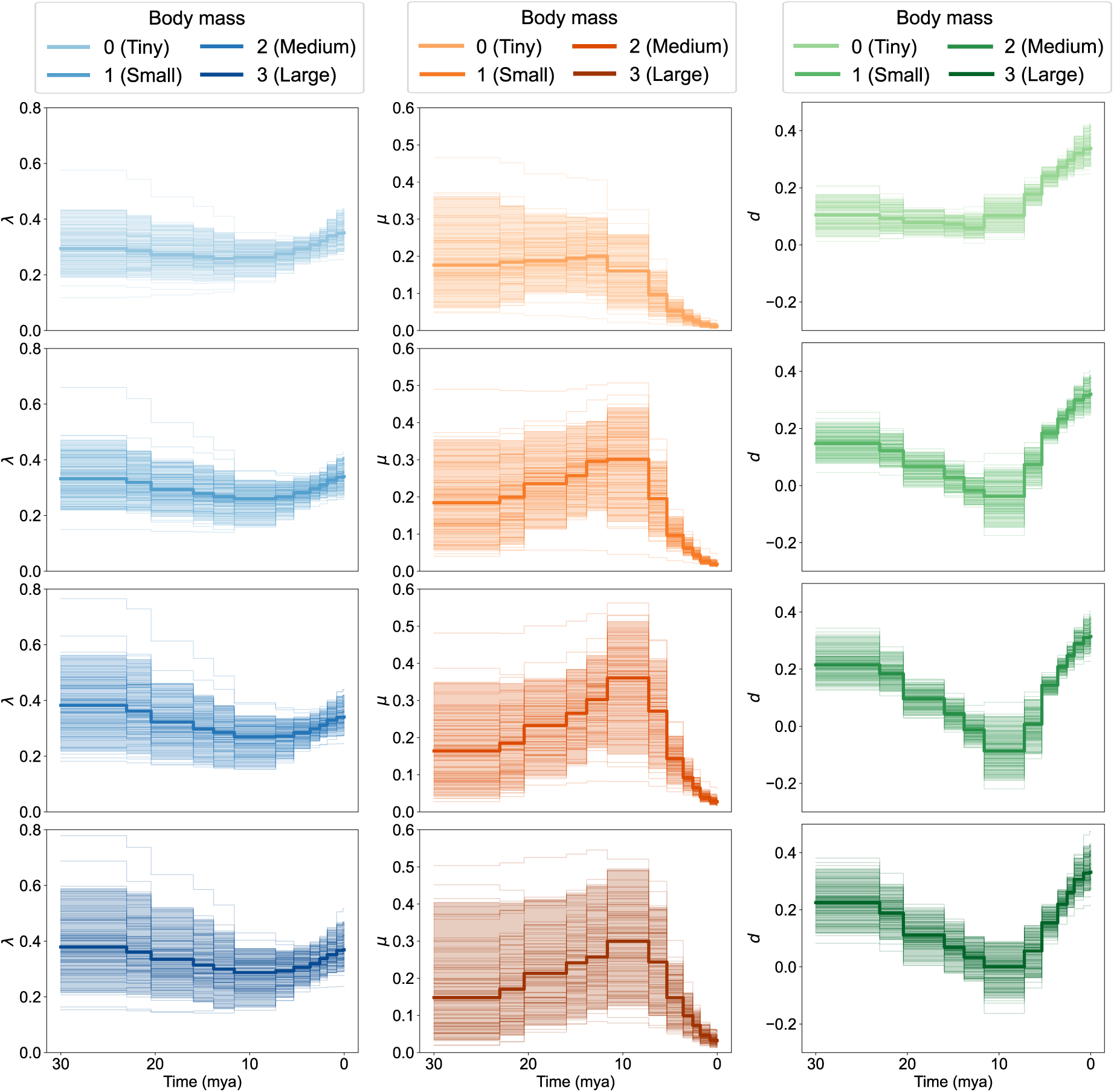
Results of BELLA for the diversification analysis of New World monkeys (Platyrrhines) phylogenies. Columns (left to right) show the speciation rate *λ*, the extinction rate *µ*, and the diversification rate *d* = *λ* − *µ*. The four rows correspond to the four body-mass categories (from smallest to largest), and display the median rate (solid line), individual phylogeny estimates (thin lines), and the 95% percentile intervals (shaded ribbons).

**Figure S5:**
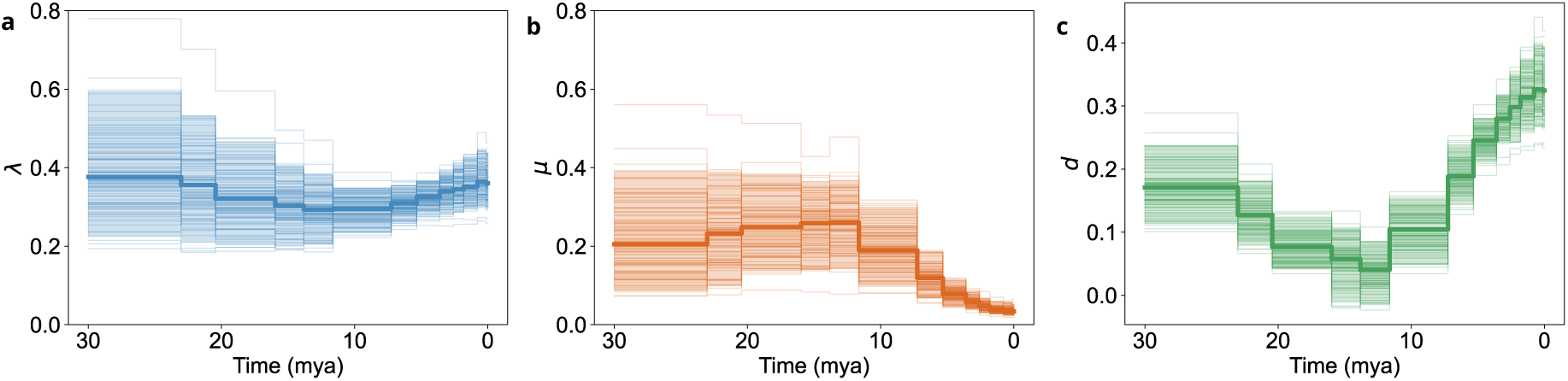
Results of BELLA for the diversification analysis of New World monkeys (Platyrrhines) phylogenies. Columns (left to right) show the marginal speciation rate *λ*, the marginal extinction rate *µ*, and the marginal diversification rate *d* = *λ* − *µ*. For each time bin, rates were averaged across body-mass classes, weighted by the number of lineages in each class. and display the median rate (solid line), individual phylogeny estimates (thin lines), and the 95% percentile intervals (shaded ribbons).

**Figure S6:**
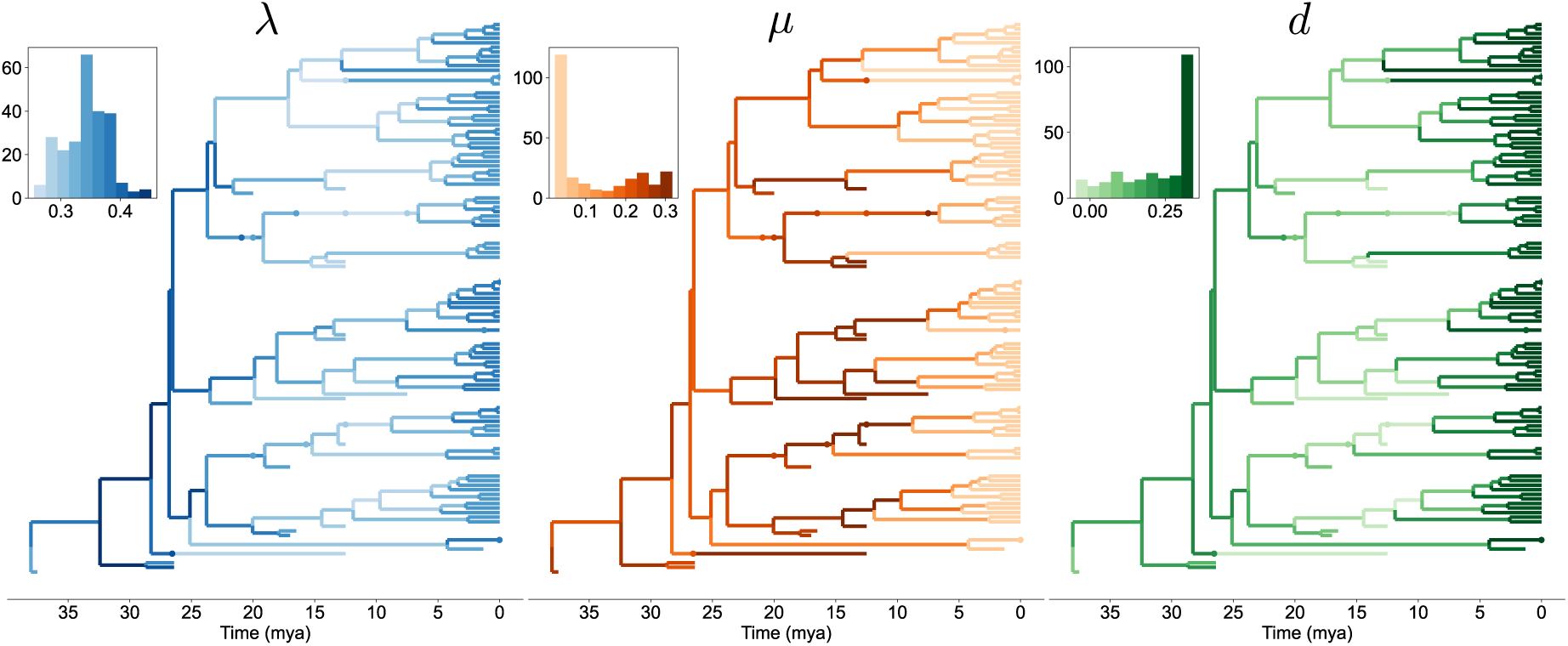
Results of BELLA for the diversification analysis of New World monkeys (Platyrrhines). From left to right, the panels show: the speciation rate (*λ*, blue), the extinction rate (*µ*, orange) and (c) the diversification rate (*d* = *λ* − *µ*, green) mapped onto the maximum clade credibility tree. Each panel includes an inset histogram illustrating the distribution of the corresponding rate across the tree.

**Figure S7:**
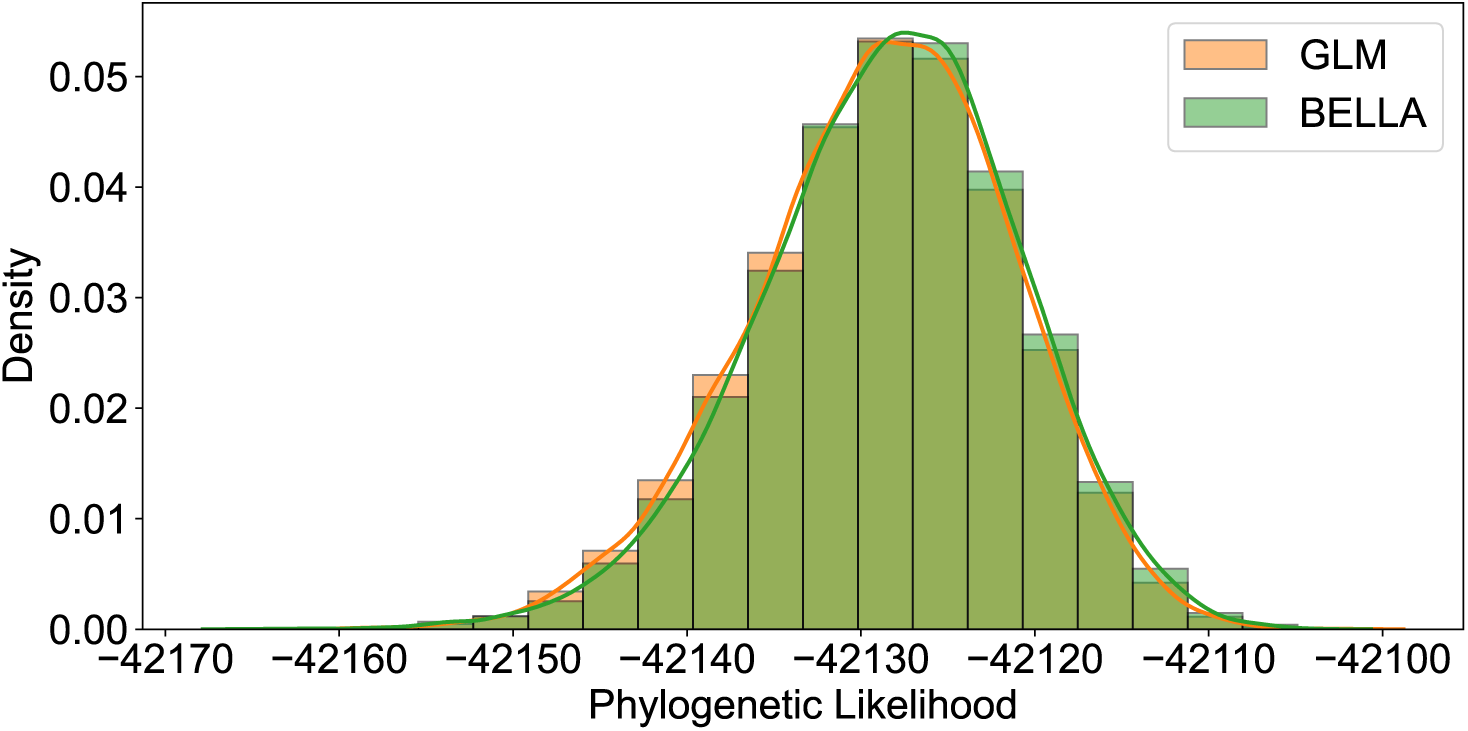
Phylogenetic likelihood distributions for the early spread of SARS-CoV-2 in Europe.

**Figure S8:**
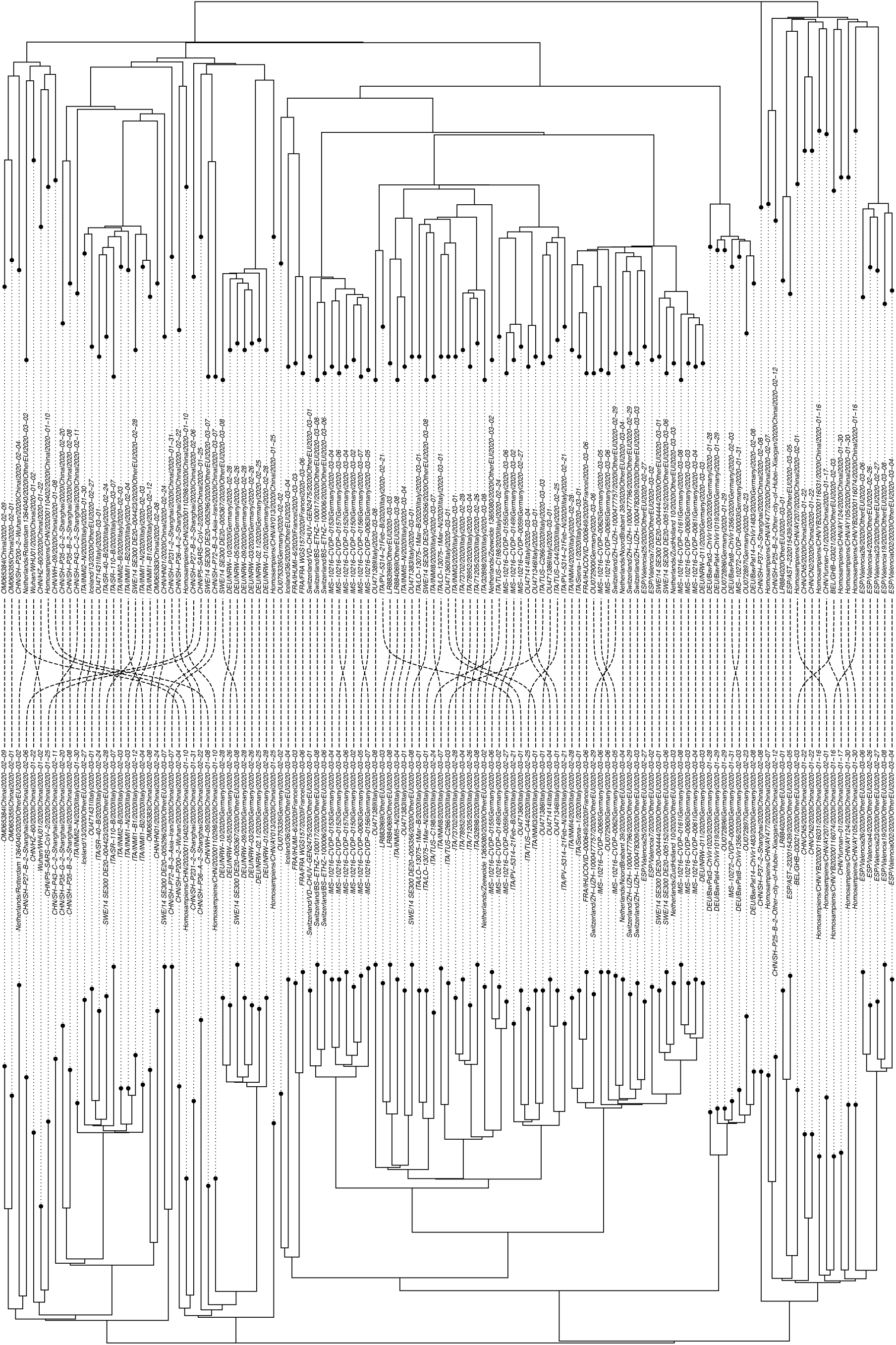
Cophylogeny comparing maximum clade credibility (MCC) trees from BELLA and GLM analyses for the early spread of SARS-CoV-2 in Europe. Lines connect identical taxa between trees.

**Figure S9:**
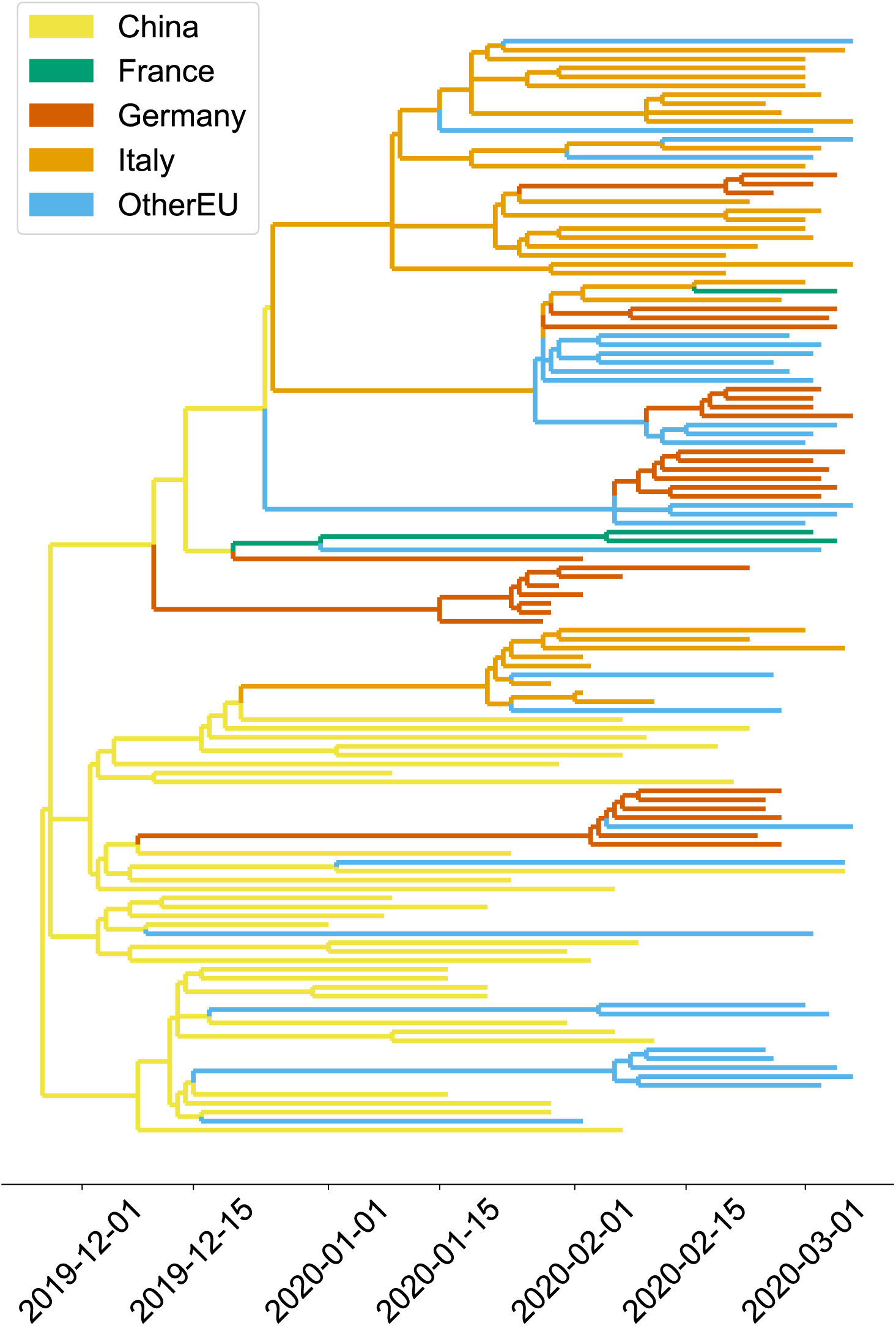
Inferred transmission tree for the early spread of SARS-CoV-2 in Europe using a GLM model.

**Figure S10:**
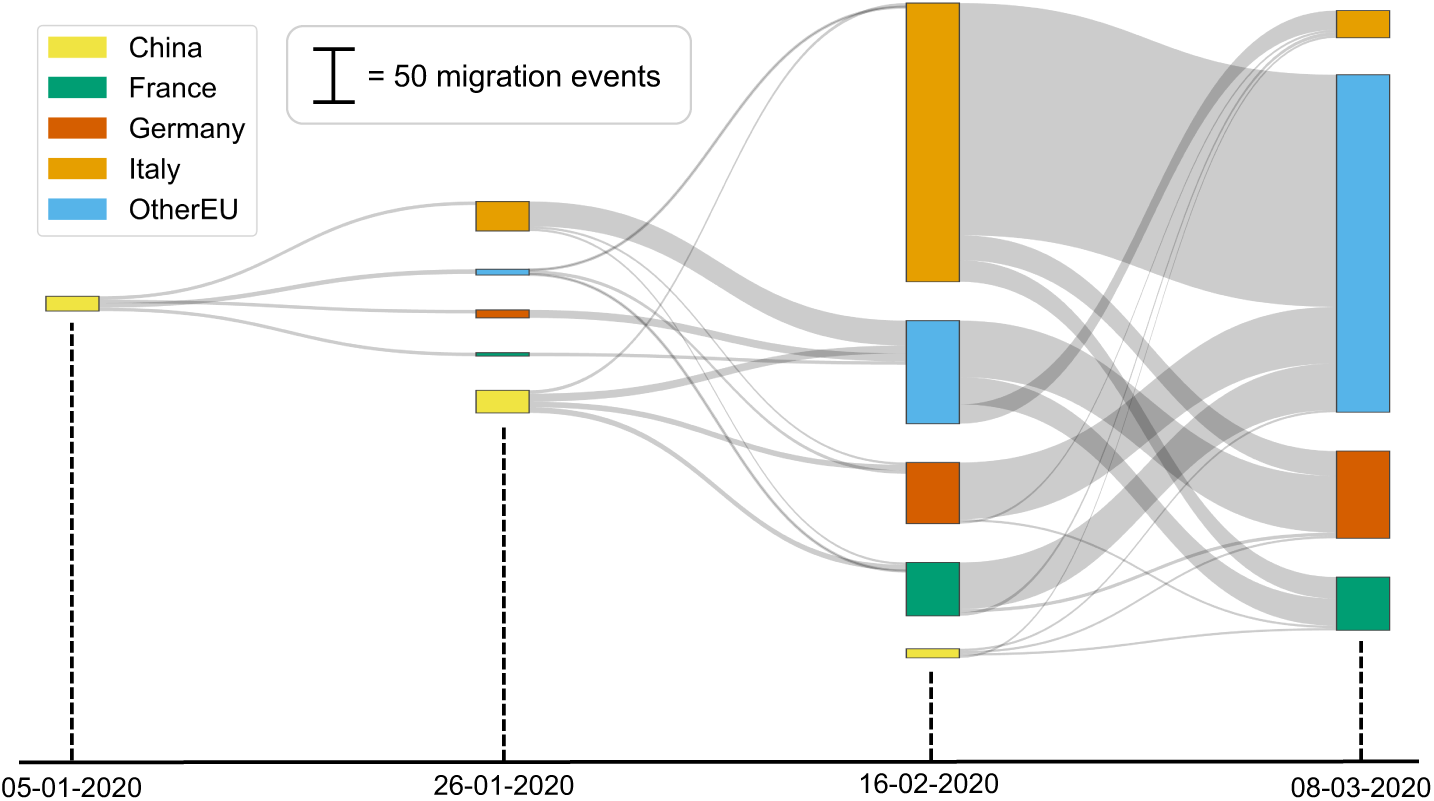
Median estimated migration fluxes between countries for the early spread of SARS-CoV-2 in Europe using a GLM model.

**Figure S11:**
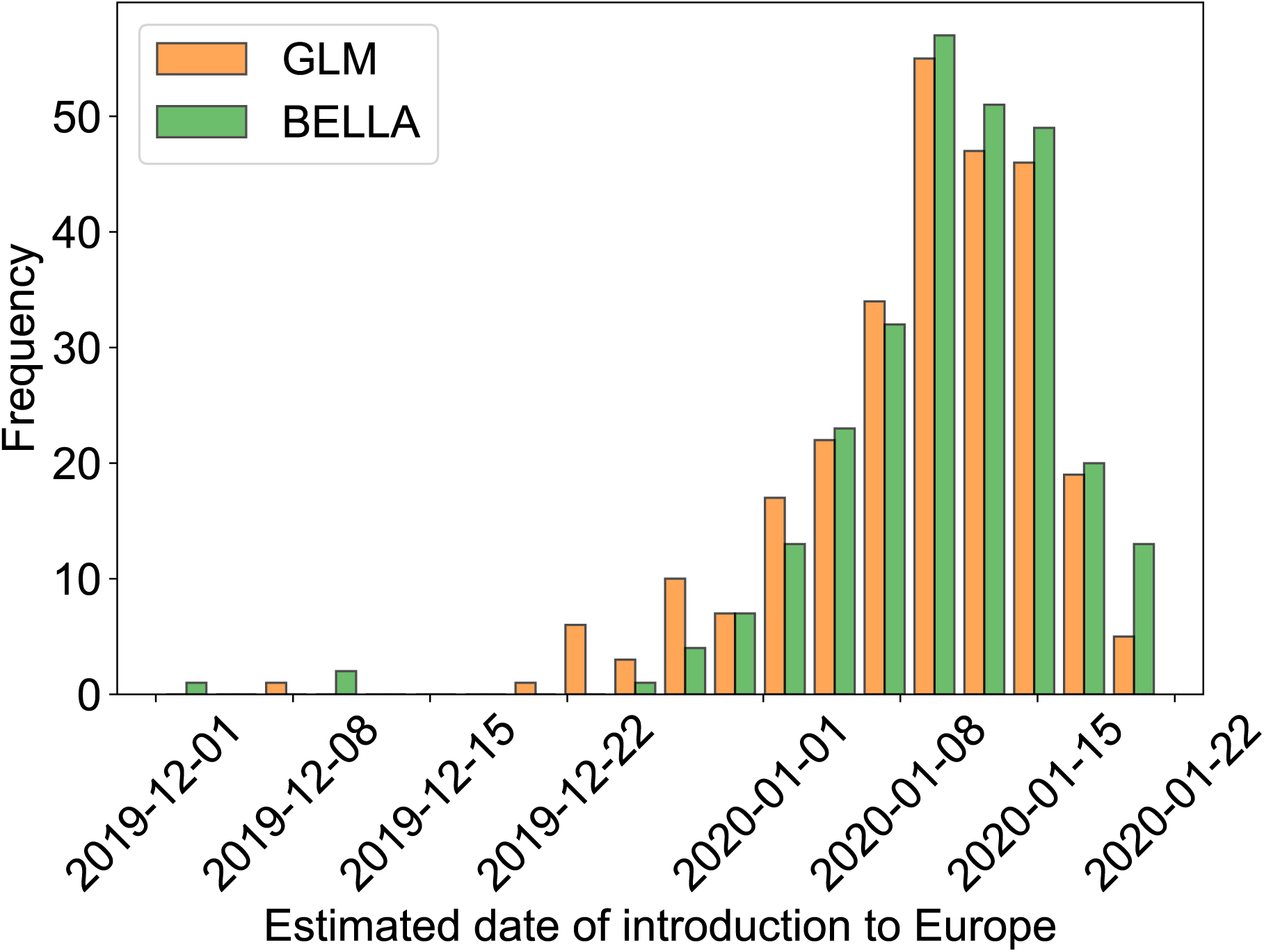
Distribution of estimated introduction dates in Europe from China during the early spread of SARS-CoV-2.

**Figure S12:**
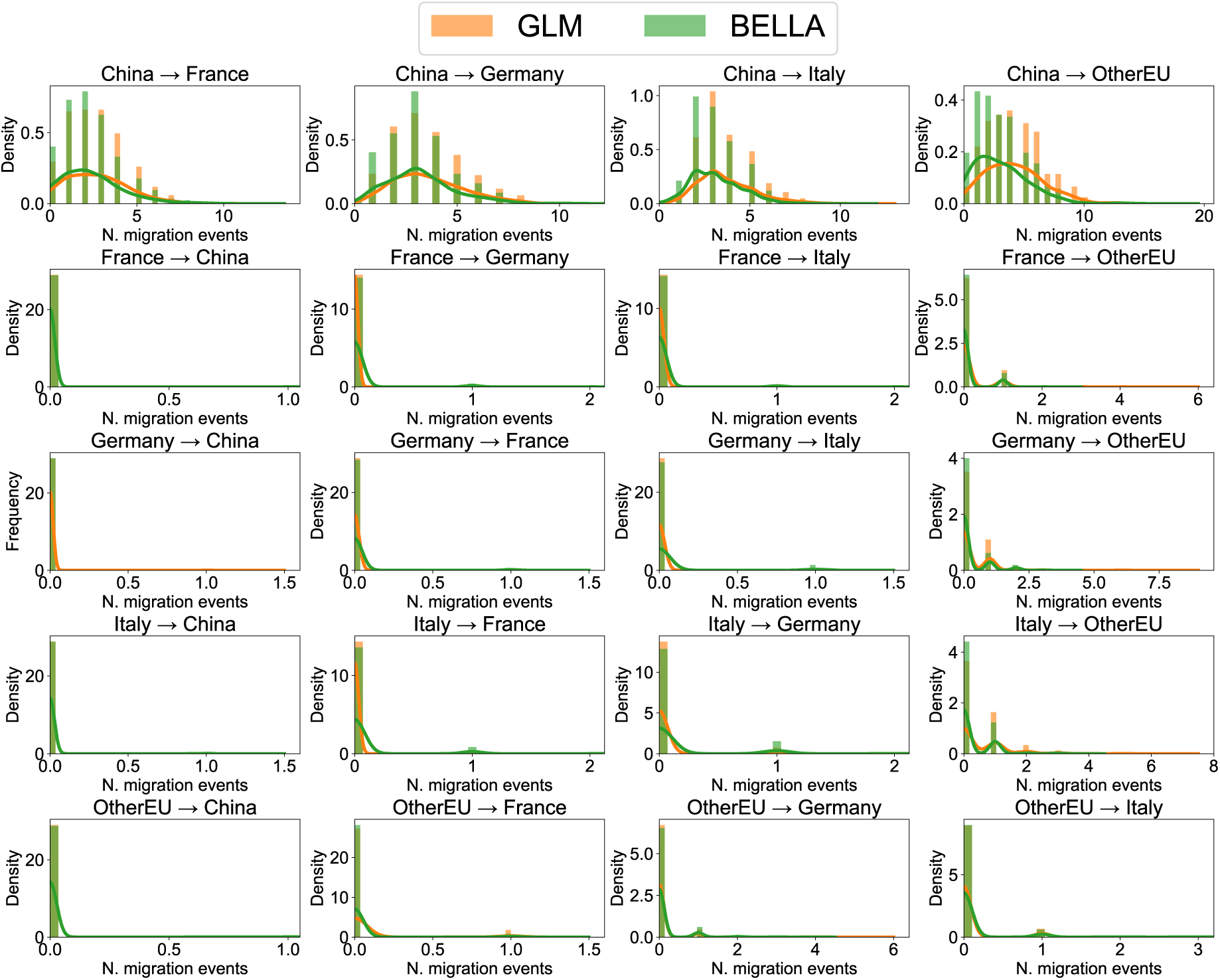
Distribution of estimated migration fluxes between countries during the early spread of SARS-CoV-2 in Europe for the period from 5 January 2020 to 26 January 2020.

**Figure S13:**
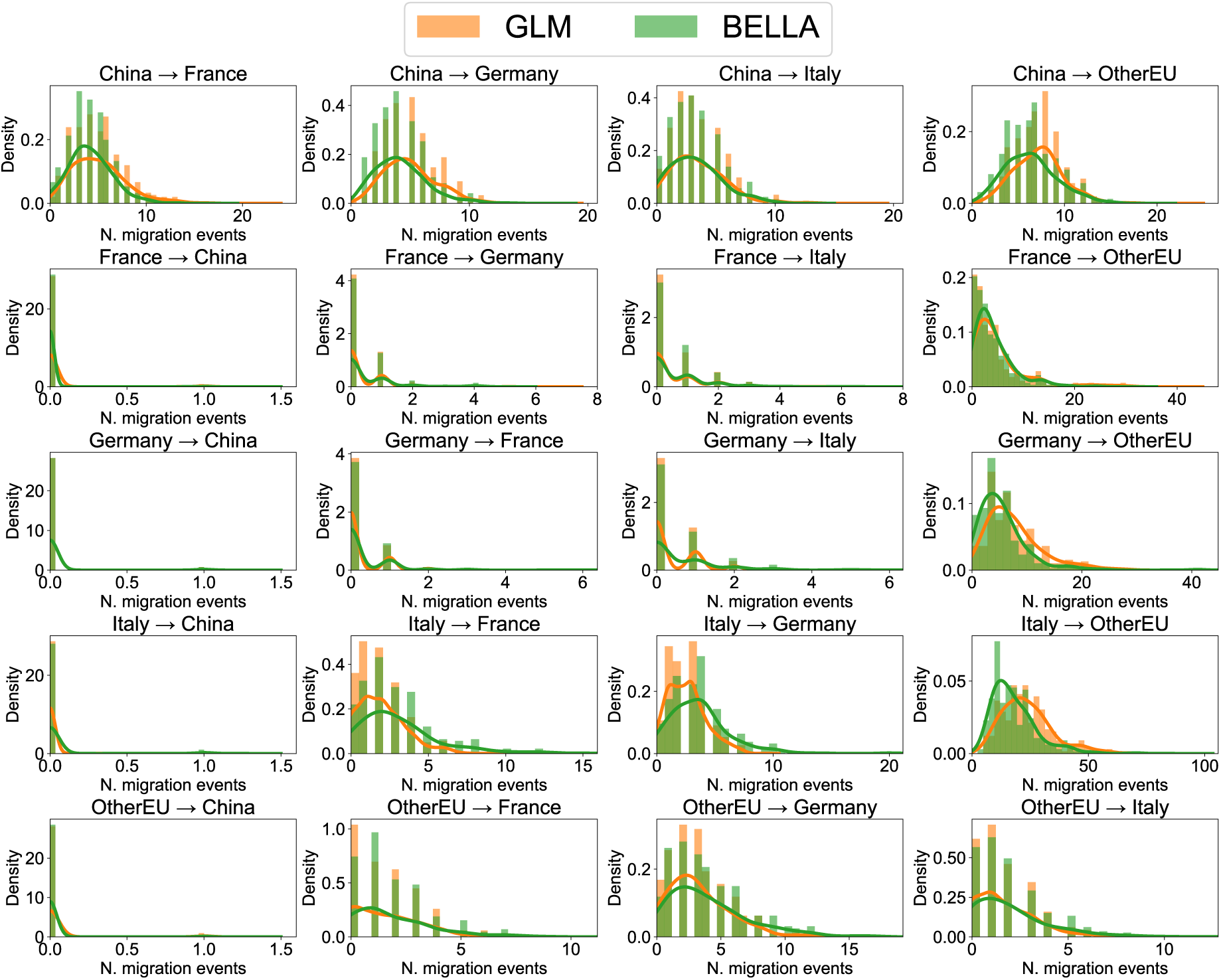
Distribution of estimated migration fluxes between countries during the early spread of SARS-CoV-2 in Europe for the period from 26 January 2020 to 16 February 2020.

**Figure S14:**
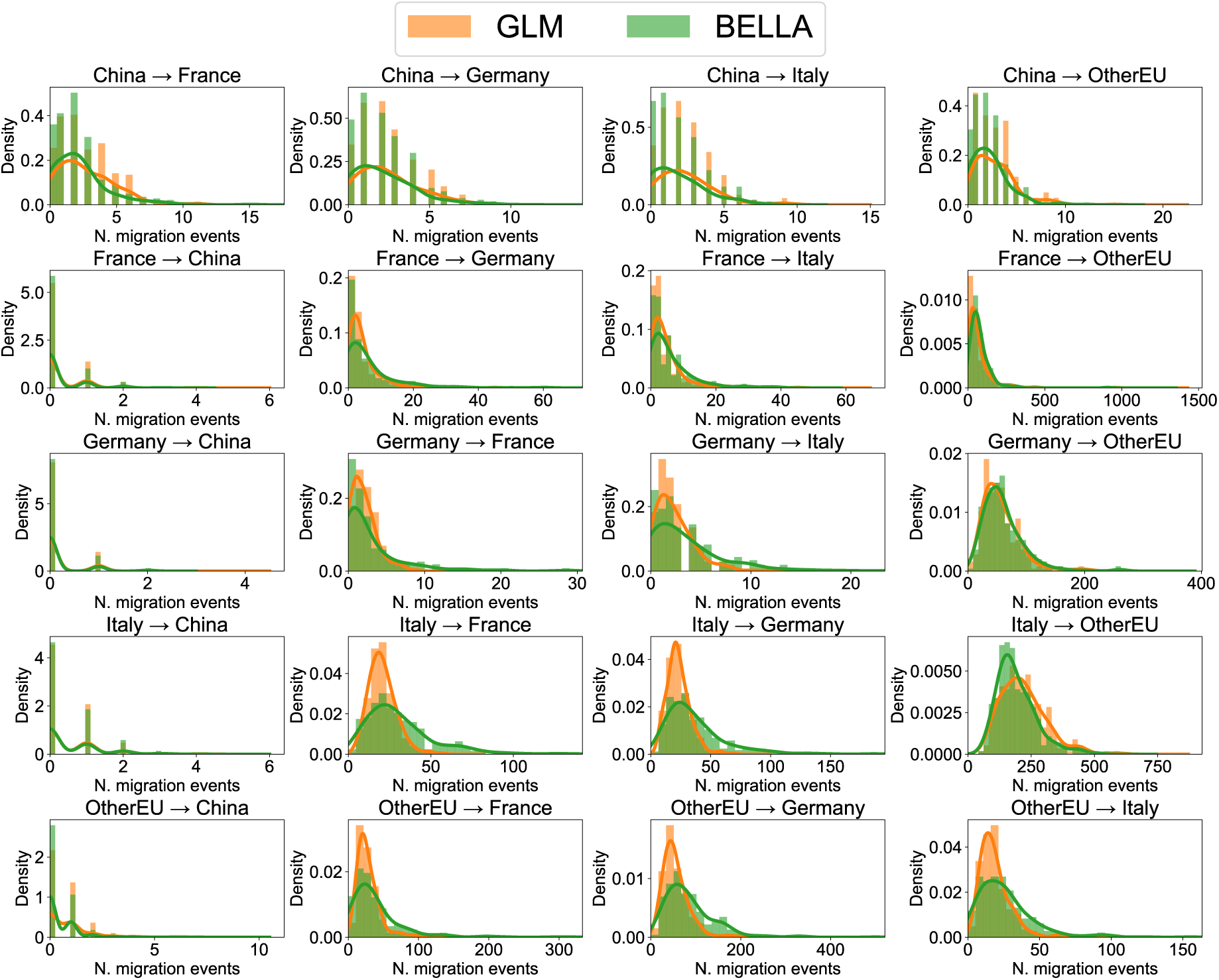
Distribution of estimated migration fluxes between countries during the early spread of SARS-CoV-2 in Europe for the period from 16 February 2020 to 8 March 2020.

### Supplementary Tables

**Table S1:** Mean absolute error (MAE) across all simulation scenarios for all inference frameworks considered (PA is the predictor-agnostic model). Bold indicates the best, underlined indicates the second-best.

| Mean absolute error (MAE) |  |  |  |  |  |  |
| --- | --- | --- | --- | --- | --- | --- |
| Scenario | Var. | PA | GLM | BELLA-3/2 | BELLA-16/8 | BELLA-32/16 |
| Epi-1 | $R_t$ | 0.109 | <u>0.052</u> | **0.032** | 0.058 | 0.065 |
| Epi-2 | $R_t$ | 0.048 | **0.006** | 0.038 | <u>0.007</u> | 0.017 |
| Epi-3 | $R_t$ | 0.055 | 0.119 | 0.084 | <u>0.054</u> | **0.036** |
| Epi-4 | $M$ | 0.058 | 0.007 | 0.005 | <u>0.004</u> | **0.003** |
| FBD-1 | $\lambda$ | 0.120 | 0.032 | **0.018** | 0.025 | <u>0.023</u> |
| | $\mu$ | 0.077 | <u>0.005</u> | **0.004** | 0.017 | 0.027 |
|  | Avg. | 0.098 | <u>0.019</u> | **0.011** | 0.021 | 0.025 |
| FBD-2 | $\lambda$ | 0.075 | 0.030 | 0.026 | <u>0.021</u> | **0.020** |
| | $\mu$ | 0.057 | 0.009 | 0.022 | **0.003** | <u>0.007</u> |
|  | Avg. | 0.066 | 0.020 | 0.024 | **0.012** | <u>0.013</u> |
| FBD-3 | $\lambda$ | 0.061 | 0.159 | **0.010** | <u>0.015</u> | 0.025 |
| | $\mu$ | 0.097 | 0.053 | **0.020** | <u>0.024</u> | 0.024 |
|  | Avg. | 0.079 | 0.106 | **0.015** | <u>0.020</u> | 0.024 |
| FBD-4 | $\lambda$ | 0.130 | 0.025 | 0.020 | <u>0.013</u> | **0.011** |
| | $\mu$ | 0.175 | 0.016 | 0.026 | <u>0.012</u> | **0.007** |
|  | Avg. | 0.153 | 0.021 | 0.023 | <u>0.013</u> | **0.009** |

**Table S2:** Coverage across all simulation scenarios for all inference frameworks considered (PA is the predictor-agnostic model).

| Coverage |  |  |  |  |  |  |
| --- | --- | --- | --- | --- | --- | --- |
| Scenario | Var. | PA | GLM | BELLA-3/2 | BELLA-16/8 | BELLA-32/16 |
| Epi-1 | $R_t$ | 0.931 | 0.964 | 0.983 | 0.990 | 0.977 |
| Epi-2 | $R_t$ | 0.954 | 0.975 | 0.979 | 0.986 | 0.984 |
| Epi-3 | $R_t$ | 0.958 | 0.748 | 0.908 | 0.975 | 0.983 |
| Epi-4 | $M$ | 0.902 | 0.606 | 0.850 | 0.961 | 0.953 |
| FBD-1 | $\lambda$ | 0.899 | 0.899 | 0.969 | 0.980 | 0.979 |
| | $\mu$ | 0.971 | 0.964 | 0.970 | 0.965 | 0.926 |
|  | Avg. | 0.935 | 0.931 | 0.969 | 0.973 | 0.952 |
| FBD-2 | $\lambda$ | 0.927 | 0.874 | 0.954 | 0.962 | 0.974 |
| | $\mu$ | 0.955 | 0.955 | 0.946 | 0.977 | 0.975 |
|  | Avg. | 0.941 | 0.914 | 0.950 | 0.970 | 0.974 |
| FBD-3 | $\lambda$ | 0.956 | 0.235 | 0.848 | 0.868 | 0.916 |
| | $\mu$ | 0.970 | 0.316 | 0.728 | 0.728 | 0.783 |
|  | Avg. | 0.963 | 0.275 | 0.788 | 0.798 | 0.850 |
| FBD-4 | $\lambda$ | 0.954 | 0.910 | 0.959 | 0.977 | 0.981 |
| | $\mu$ | 0.892 | 0.903 | 0.923 | 0.976 | 0.968 |
|  | Avg. | 0.923 | 0.907 | 0.941 | 0.977 | 0.974 |

**Table S3:** Average 95% credible interval width across all simulation scenarios for all inference frameworks considered (PA is the predictor-agnostic model).

| Average 95% credible interval width |  |  |  |  |  |  |
| --- | --- | --- | --- | --- | --- | --- |
| Scenario | Var. | PA | GLM | BELLA-3/2 | BELLA-16/8 | BELLA-32/16 |
| Epi-1 | $R_t$ | 0.989 | 0.373 | 0.307 | 0.537 | 0.686 |
| Epi-2 | $R_t$ | 0.931 | 0.354 | 0.380 | 0.523 | 0.659 |
| Epi-3 | $R_t$ | 1.131 | 0.435 | 0.460 | 0.642 | 0.778 |
| Epi-4 | $M$ | 0.163 | 0.034 | 0.035 | 0.051 | 0.067 |
| FBD-1 | $\lambda$ | 0.460 | 0.146 | 0.117 | 0.185 | 0.228 |
| | $\mu$ | 0.461 | 0.120 | 0.096 | 0.135 | 0.157 |
|  | Avg. | 0.460 | 0.133 | 0.106 | 0.160 | 0.193 |
| FBD-2 | $\lambda$ | 0.357 | 0.139 | 0.147 | 0.185 | 0.218 |
| | $\mu$ | 0.389 | 0.110 | 0.104 | 0.148 | 0.179 |
|  | Avg. | 0.373 | 0.124 | 0.126 | 0.166 | 0.199 |
| FBD-3 | $\lambda$ | 0.403 | 0.174 | 0.165 | 0.213 | 0.227 |
| | $\mu$ | 0.398 | 0.094 | 0.093 | 0.106 | 0.118 |
|  | Avg. | 0.400 | 0.134 | 0.129 | 0.159 | 0.173 |
| FBD-4 | $\lambda$ | 0.708 | 0.176 | 0.203 | 0.265 | 0.313 |
| | $\mu$ | 0.818 | 0.151 | 0.182 | 0.251 | 0.318 |
|  | Avg. | 0.763 | 0.163 | 0.193 | 0.258 | 0.315 |

**Table S4:**
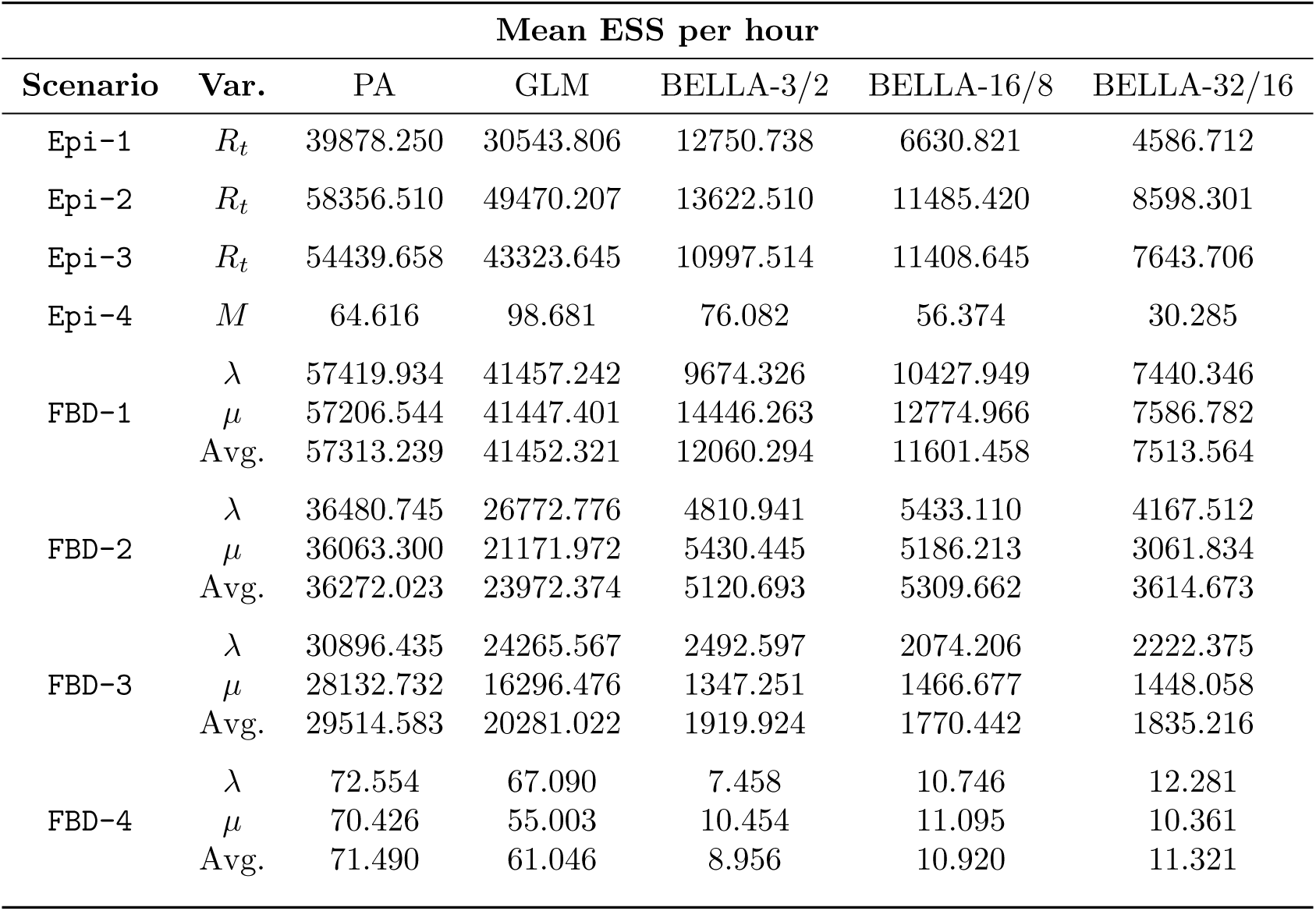
Mean effective sample size (ESS) per hour across all simulation scenarios for all inference frameworks considered (PA is the predictor-agnostic model).

